# ZNF574 is a Quality Control Factor For Defective Ribosome Biogenesis Intermediates

**DOI:** 10.1101/2024.04.26.591394

**Authors:** Jared F. Akers, Adrian Bothe, Hanna Suh, Carmen Jung, Zachary Stolp, Tanushree Ghosh, Liewei Yan, Yuming Wang, Tarabryn Grismer, Andreas Reyes, Tianyi Hu, Shouling Xu, Nenad Ban, Kamena K. Kostova

**Affiliations:** Carnegie Institution for Science, Biosphere Science and Engineering Division, Baltimore, MD, 21218, USA; Department of Biology, Institute of Molecular Biology and Biophysics, ETH Zurich, 8093 Zurich, Switzerland

## Abstract

Eukaryotic ribosome assembly is an intricate process that involves four ribosomal RNAs, 80 ribosomal proteins, and over 200 biogenesis factors that take part in numerous interdependent steps. This complexity creates a large genetic space in which pathogenic mutations can occur. “Dead-end” ribosome intermediates that result from biogenesis errors are rapidly degraded, affirming the existence of quality control pathway(s) that monitor ribosome assembly. However, the factors that differentiate between on-path and dead-end intermediates are unknown. We engineered a system to perturb ribosome assembly in human cells and discovered that faulty ribosomes are degraded via the ubiquitin proteasome system. We identified *ZNF574* as a key component of a novel quality control pathway, which we term the <u>R</u>ibosome <u>A</u>ssembly <u>S</u>urveillance <u>P</u>athway (RASP). Loss of ZNF574 results in the accumulation of faulty biogenesis intermediates that interfere with global ribosome production, further emphasizing the role of RASP in protein homeostasis and cellular health.

## Introduction

The eukaryotic ribosome is a complex molecular machine responsible for the translation of mRNA to protein. It contains both RNA and protein, and its intricate assembly is facilitated by approximately 200 protein factors, and numerous small RNAs^1^. These components engage in hundreds of spatially and temporally coordinated reactions spanning the nucleolus, nucleoplasm, and cytoplasm^2,3^. This delicate process is susceptible to various forms of disruption and damage. For instance, environmental stresses like ultraviolet radiation^4–7^ and reactive oxygen species^8,9^ can damage both the ribosomal RNA (rRNA) and ribosomal proteins. Genetic perturbations^10,11^, errors during transcription^12,13^ and translation^14,15^, as well as compromised post-transcriptional modifications^16^ can impair ribosome biogenesis, yielding non-functional biogenesis intermediates. Faulty ribosome production leads to decreased protein synthesis, increased cellular stress, and often cell death^17^. On the organismal level, ribosome biogenesis defects manifest as diverse pathological phenotypes across tissues and developmental stages, such as severe anemias, craniofacial abnormalities, growth delays, intellectual disabilities, and cancer predisposition^18–20^. If left unresolved, accumulation of defective ribosomes could have dire consequences for cell viability and human health.

Previous research on ribosome biogenesis primarily focused on factors facilitating the proper assembly of the ribosome^21–23^. By contrast, investigations into quality control mechanisms targeting abnormal pre-ribosomes remain relatively unexplored. Nonetheless, a limited number of studies hint at the existence of quality control pathways capable of detecting and eliminating defective ribosomes. For instance, in plants, UV-B radiation induces crosslinking between the ribosomal proteins and the ribosomal RNA, inhibiting protein production^5–7^. However, removal of the damaging agent allows for clearance of the defective ribosomes, restoring translation to normal levels. Similarly, studies in yeast have shown that ribosomes containing a mutated rRNA are targeted by a dedicated quality control pathway, termed nonfunctional rRNA decay (NRD)^24,25^. Furthermore, experiments in yeast and cell line models of defective ribosome assembly^18,20,26^ have shown rapid degradation of faulty biogenesis intermediates. Despite the compelling evidence that quality control pathways exist, our understanding of the factors involved and their mechanism of identifying and degrading faulty ribosomes remains limited.

Studying ribosome quality control has been hindered by the lack of robust methods for identifying and isolating defective ribosomes. Although there are no studies measuring the fidelity of ribosome assembly, we can do a back-of-the-envelope calculation for the fraction of defective ribosomes for HeLa cells, which assemble approximately 7,500 ribosomes per minute^27^. Assuming the error rate for RNA polymerase is 10^-6^ errors per nucleotide, the total length of the human ribosomal RNA is 7216 nucleotides, and the conservation of the ribosomal RNA sequence between yeast and human is approximately 60%, implying that these residues are intolerant to mutational changes, then HeLa cells are expected to assemble ∼30 defective ribosomes per minute. Given that HeLa cells typically contain about 3 million ribosomes^28^, detecting this small fraction of endogenously produced defective ribosomes is akin to finding a needle in a haystack. Attempts have been made to increase the number of faulty ribosomes by subjecting cells to damaging agents, such as UV^4,29^ or oxidative stress^30^, or by mutating or deleting assembly factors to block ribosome biogenesis^31^. However, such conditions severely impact cell viability since protein production is essential for cellular functions, complicating the interpretation of results due to secondary effects^32–34^.

To overcome these challenges, we established a method to impede the maturation of a small subset of ribosomes in human cells. The assembly of the 60S large ribosomal subunit begins in the nucleolus, continues through the nucleoplasm, and concludes in the cytoplasm with the incorporation of the ribosomal protein uL16/RPL10^35^ and the release of the late-acting biogenesis factors^36^. We designed a system where incorporation of a mutant uL16 lacking residues 102-111 blocks the assembly of the maturing large subunit (Fig 1I). This approach offers several advantages over methods relying on mutations in ribosome biogenesis factors to induce ribosome assembly failure. First, we can specifically monitor and isolate defective biogenesis intermediates from the global pool of ribosomes by introducing epitope tags on uL16^mut^. Second, since uL16^mut^ is ectopically expressed at low levels, only a fraction of the large subunits fails to assemble, allowing for continuous ribosome production and cell viability. Finally, by strategically mutating uL16, we can block the final step of ribosome maturation, enabling us to utilize the extensive structural^37–39^ and biochemical^40–42^ data on productive ribosome assembly to compare to ribosome biogenesis failure.

**Fig. 1.**
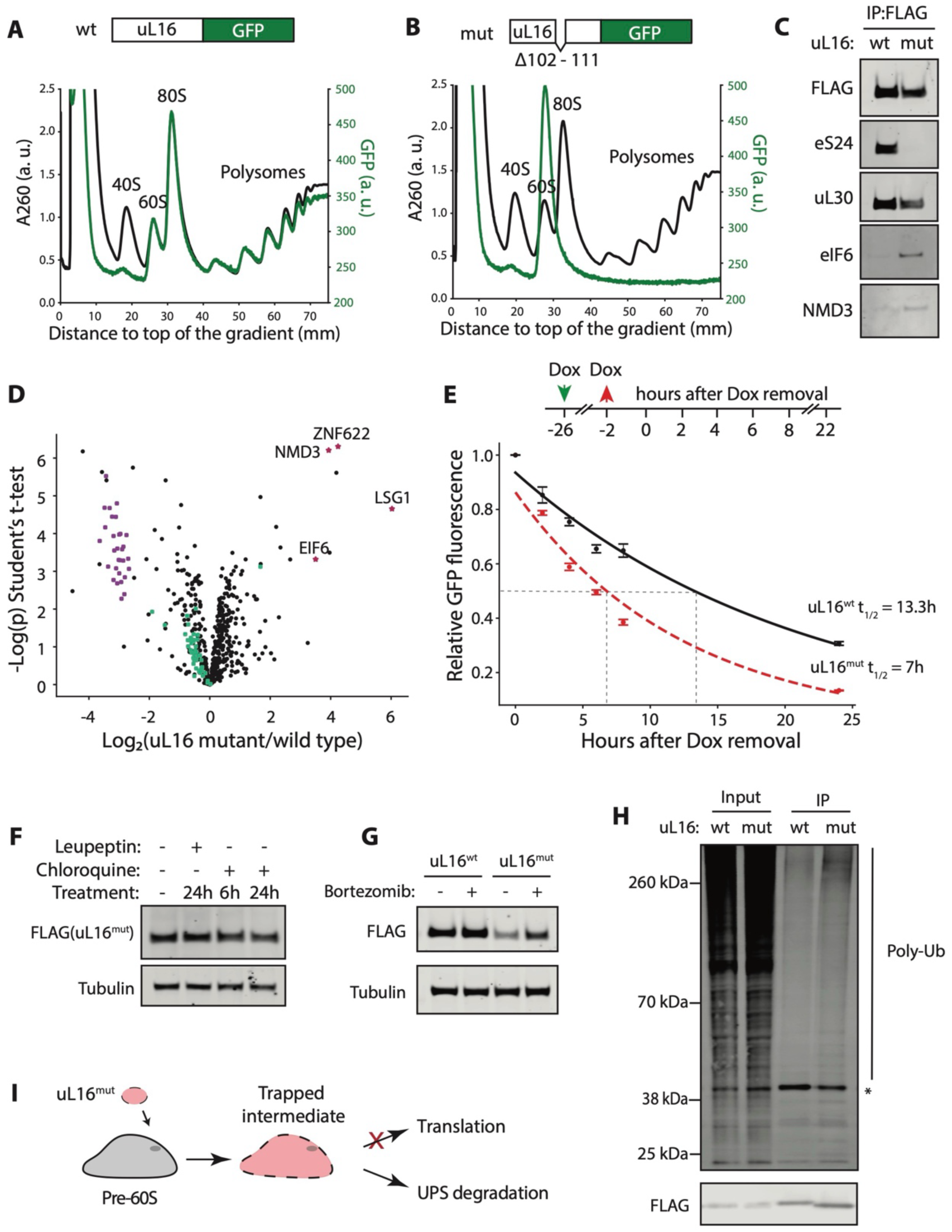
Incorporation of uL16^mut^ blocks maturation of the ribosomal large subunit. **A, B.** Top: diagrams of wild type and mutant uL16-GFP constructs. Bottom: Polysome gradients from K562 cells transduced with GFP-tagged wild type (A) or mutant (B) uL16. The total RNA absorbance at A260 nm (black line) and GFP signal (green line) were measured along the gradients. **C, D.** FLAG-tagged wild type or mutant uL16 was immunoprecipitated and the associated proteins were identified via immunoblotting (C) or mass spectrometry (D). Mass spectrometry data was analyzed to identify the fold-enrichment of proteins immunoprecipitated with uL16^mut^ compared with uL16^wt^. Highlighted are ribosomal proteins from the small subunit (purple squares), large subunit (green squares), and biogenesis factors (red stars). **E.** Wild type or mutant uL16-GFP was expressed under a doxycycline (Dox)-inducible promoter. Cells were treated with Dox for 24 hours, the drug was washed away, and the levels of GFP-tagged uL16 were measured over time via flow cytometry (timeline of the experiment shown on top). Half-lives were calculated by fitting an exponential decay curve (mean ± SD, N=3). **F.** K562 cells expressing ul16^mut^ were treated with the autophagy inhibitors leupeptin and chloroquine. uL16 levels were measured via western blot. Tubulin was used as loading control. **G.** K562 cells expressing wild type or mutant uL16 were treated with the proteasome inhibitor Bortezomib and uL16 levels were measured via western blot. Tubulin was used as loading control. **H.** FLAG-tagged wild type or mutant uL16 was immunoprecipitated and the presence of poly ubiquitin (poly-Ub) was detected via western blot. Non-specific bands are labeled with an asterisk (*). **I.** Diagram of uL16^mut^ incorporation into maturing large pre-60S subunits. These subunits cannot complete assembly and become dead-end intermediates that are degraded via the ubiquitin-proteasome system (UPS).

Leveraging this system, we determined that defective ribosome biogenesis intermediates are rapidly degraded in human cells. Through CRISPR interference (CRISPRi) screens, we identified *ZNF574* as a key component of a novel quality control pathway, which we term the <u>R</u>ibosome <u>A</u>ssembly <u>S</u>urveillance <u>P</u>athway (RASP). Disruption of RASP leads to the accumulation of defective biogenesis intermediates that ultimately hinder the assembly of new ribosomes, further emphasizing the critical role of ribosome quality control in protein homeostasis.

## Results

### Modeling ribosome biogenesis failure in human cells

To investigate the mechanisms underlying the detection and elimination of defective ribosome biogenesis intermediates, we engineered a human cell line where a subset of large subunits fails to fully mature upon incorporation of ribosomal protein uL16 (uL16^mut^), an integral component of the large subunit. The mutant lacks residues 102-111, which are critical for the formation of the peptidyl transferase center^43^. Expression of uL16^mut^ in yeast results in impaired translation, despite its structural incorporation into the ribosome^41^. This observation led to a model where large subunits harboring uL16^mut^ are trapped during the late stages of ribosome assembly and cannot engage in active translation.

Previous studies in yeast have shown that cells solely expressing uL16^mut^ are not viable^41^. We therefore ectopically expressed tagged wild type uL16 (uL16^wt^) or uL16^mut^ at low levels in human cells (SFig. 1). As a result, the majority of the maturing large subunits incorporate the endogenous wild type uL16, ensuring the production of a sufficient number of functional ribosomes to sustain cellular health. This system gave us an opportunity to study dead-end ribosome biogenesis intermediates without severely compromising cellular health.

To examine the uL16^mut^ ribosome biogenesis phenotype in human cells, we ectopically expressed a GFP-tagged uL16^wt^ or uL16^mut^ in K562 cells and fractionated the ribosome populations using sucrose gradients (Fig. 1A, B). The assembly state and translational status of all ribosomes were measured using A_260_ absorption and ribosomes containing GFP tagged uL16 were monitored using GFP fluorescence. uL16^wt^-GFP co-migrated with 60S large subunits, 80S monosomes, and translating polysomes (Fig. 1A), confirming that the GFP tag does not interfere with ribosome assembly or active translation. The minor peak detected in the 40S fraction is non-specific and can be seen in sucrose gradients from K562 cells lacking GFP expression (SFig. 2). In contrast, the uL16^mut^-GFP co-sedimented exclusively with the 60S peak. This observation confirms the incorporation of the mutant protein into 60S subunits (60S^mut^), however, these large subunits are unable to engage in active translation. Similar results were observed with FLAG-tagged uL16 (SFig. 3).

To confirm that uL16^mut^ incorporation blocks 60S maturation, we purified tagged uL16^wt^ or uL16^mut^ and identified associated proteins via western blotting (Fig. 1C) and mass spectrometry analysis (Fig. 1D). We did not detect an interaction between uL16^mut^ and small subunit proteins, validating that the 60S^mut^ subunits are trapped during biogenesis and unable to pair with small subunits to engage in active translation. We also observed significant enrichment of late ribosome biogenesis factors, such as eIF6^44^ and NMD3^40,42,45^, upon uL16^mut^ immunoprecipitation. eIF6 functions as an anti-association factor that prevents premature joining of 40S subunits^44^, thereby providing a mechanistic explanation for the failure of 60S^mut^ to associate with ribosomal proteins from the small subunit.

### 60Smut subunits are rapidly degraded during biogenesis via the ubiquitin-proteasome system, and not autophagy

Faulty ribosomes are rapidly detected and cleared in eukaryotic cells^46^. For example, ribosomes harboring mutations in key rRNA residues are promptly degraded via the non-functional rRNA decay (NRD) pathways^24,25^. Furthermore, depletion in ribosome synthesis factors that interfere with rRNA processing also triggers the stabilization of non-canonical biogenesis intermediates^47^. We hypothesized that pre-60S ribosomes containing uL16^mut^ are also subjected to quality control and degradation. To measure the half-life of GFP-tagged uL16^wt^ or uL16^mut^, we induced the expression of these proteins under a doxycycline-inducible promoter^48^ in K562 cells. Following a 24-hour pulse of doxycycline, we stopped uL16-GFP synthesis by removal of the drug and monitored the decrease in the protein levels over time via flow cytometry (Fig. 1E, SFig. 4). Our analysis revealed a half-life of approximately 13 hours for uL16^wt^, whereas uL16^mut^ exhibited a shorter half-life of approximately 7 hours, confirming the active degradation of the mutant protein.

Next, we determined whether the uL16^mut^-containing large subunits are targeted for degradation during biogenesis or active translation in the cytoplasm. Although we did not detect uL16^mut^ co-migrating with actively translating ribosomes on polysome gradients (Fig. 1B), it is possible that a small fraction of the defective large subunits may engage mRNAs, but fail to translate, causing ribosome collisions^49^ and subsequent ribosome quality control mechanisms^50^. To explore this possibility, we performed a pulse-chase experiment in the presence of the elongation inhibitor cycloheximide^51^ (SFig. 5). If the degradation of uL16^mut^-containing ribosomes is triggered by active translation, blocking elongation with cycloheximide will interfere with the detection of faulty large subunits. However, cycloheximide treatment did not prolong the half-life of uL16^mut^, confirming that these faulty large subunits are targeted for degradation during biogenesis and not active translation.

In eukaryotic cells, ribosome degradation occurs through two major pathways: autophagy^52^ and cytosolic turnover of ribosomal proteins by the proteasome^53–55^. Treating the cells with the autophagy inhibitors leupeptin^56^ or chloroquine^57^ did not stabilize uL16^mut^ (Fig. 1F). To directly measure uL16 targeting to the lysosome, we transduced K562 cells with uL16^wt^ or uL16^mut^ tagged with the pH sensitive fluorophore Keima^58,59^. Tagging uL16 with Keima did not interfere with ribosome assembly or active translation (SFig. 6A, B). Lysosomal targeting of uL16^mut^ would result in a higher pH5/pH7 Keima signal, similar to Torin1-induced autophagy^60^ (SFig. 6C). However, uL16^mut^-Keima did not exhibit a higher pH5/pH7 signal compared to the uL16^wt^. Together, these results suggest that uL16^mut^ is not degraded via autophagy.

To probe the role of the ubiquitin-proteasome system (UPS) in uL16^mut^ degradation, we treated K562 cells expressing FLAG-tagged uL16^wt^ or uL16^mut^ with the proteasome inhibitor bortezomib^61^ (Fig. 1G). We observed a stabilization of uL16^mut^, but not uL16^wt^, upon inhibition of UPS degradation. Further, we detected polyubiquitination in immunoprecipitated samples of uL16^mut^ (Fig. 1H), confirming that the trapped biogenesis intermediates obtained by uL16^mut^ incorporation are degraded via the ubiquitin-proteasome system (Fig. 1I).

### CRISPRi screens identify ZNF574 as a key <u>R</u>ibosome <u>A</u>ssembly <u>S</u>urveillance <u>P</u>athway (RASP) factors

Several quality control factors are known to interact with mature ribosomes^50,53,54^ or individual ribosomal proteins^62–65^. To test if these factors have a role in uL16^mut^ degradation, we systematically knocked them down using CRISPR interference (CRISPRi)^66^ in K562 cells expressing a uL16^mut^-RFP reporter or a control uL16^wt^-RFP construct (Fig. 2A). None of the knockdowns led to stabilization of uL16^mut^-RFP, suggesting that previously characterized quality control pathways do not target uL16^mut^-containing 60S subunits. Therefore, we concluded that 60S^mut^ subunits are cleared by a dedicated quality control pathway that we termed RASP (<u>R</u>ibosome <u>A</u>ssembly <u>S</u>urveillance <u>P</u>athway).

**Fig. 2.**
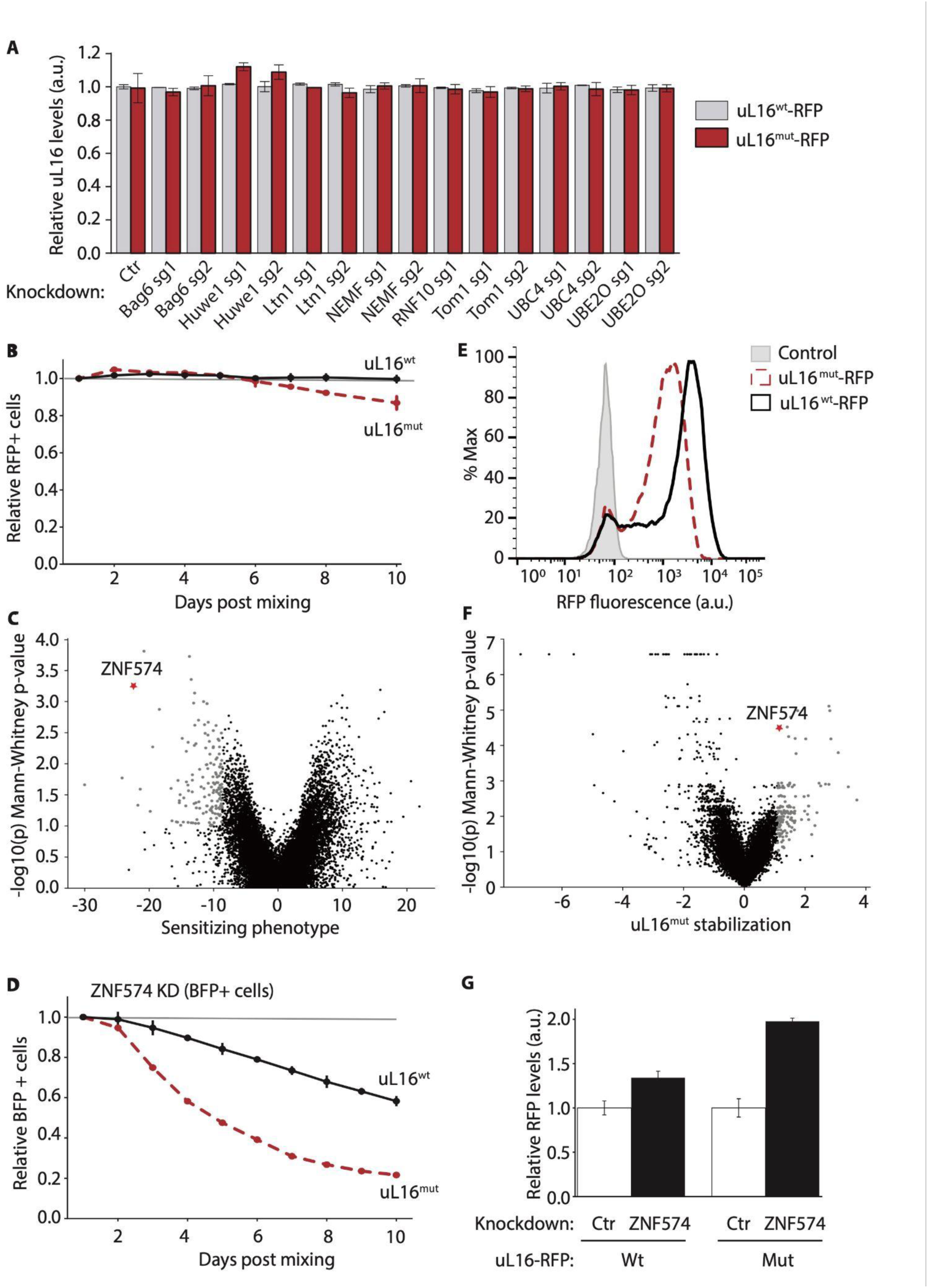
CRISPRi screens for ribosome biogenesis quality control factors. **A.** Known quality control factors were knocked down via CRISPRi and the levels of wild type and mutant uL16 were measured via flow cytometry (mean ± SD, N=3). Values are normalized to control knockdown (Ctr). **B.** K562 cells expressing RFP-tagged wild type or mutant uL16 were mixed with control cells and the ratio of RFP positive to RFP negative cells was measured at several timepoints (mean ± SD, N=3). **C.** Growth-based CRISPRi screen for quality control factors targeting uL16^mut^. The sensitizing phenotype was calculated by comparing the sgRNA abundance between cells expressing wild type or mutant uL16 after 10 cell doublings. Highlighted in gray are the top 150 hits from the screen. ZNF574 is highlighted with a red asterisk. Metrics are defined quantitatively in STAR Methods. **D.** K562 cells expressing wild type or mutant uL16 were transduced with BFP marked sgRNA targeting ZNF574. The ratio of ZNF574 knockdown to control cells (BFP positive vs BFP negative) was measured at several timepoints (mean ± SD, N=3). **E.** Protein levels of RFP-tagged wild type of mutant uL16 were measured via flow cytometry. Control cells represent the basal autofluorescence of the K562 cell line. **F.** FACS-based CRISPRi screen for quality control factors targeting uL16^mut^. The uL16^mut^ stabilization was calculated by comparing the sgRNA abundance between cells with high versus low uL16^mut^-RFP levels. Highlighted in gray are the top 150 hits from the screen. ZNF574 is highlighted in red. Metrics are defined quantitatively in STAR Methods. **G.** Protein levels of wild type or mutant uL16 were measured upon control or ZNF574 knockdown via flow cytometry (mean ± SD, N=3).

To identify novel RASP factors, we performed two independent CRISPRi screens measuring different phenotypes^67,68^. First, we investigated whether cells expressing uL16^mut^ exhibited a growth defect relative to uL16^wt^. We mixed control K562 cells with K562 cells expressing wild type or mutant uL16-RFP and measured the proportion of RFP positive cells over time via flow cytometry (Fig. 2B). Cells expressing uL16^mut^-RFP were slowly outcompeted by wild type cells, suggesting that although these cells were actively degrading the uL16^mut^-containing large subunits, the presence of faulty biogenesis intermediates compromised cellular fitness. We then hypothesized that RASP inhibition will further impede the proliferation rate of uL16^mut^ cells compared to those expressing uL16^wt^ and designed a growth based CRISPRi screen to detect this phenotype (SFig. 7A). We engineered two cell lines: one expressing uL16^mut^ and a control cell line expressing uL16^wt^, both constitutively expressing the dCas9-KRAB CRISPRi effector. We transduced the two cell lines with the sgRNA library (hCRISPRi-v2)^67^ targeting all known human protein-coding genes (SFig. 7A) and compared the change in sgRNA abundance after ten cell doublings between the two cell lines (Fig. 2C). We expected sgRNAs targeting RASP factors to be depleted in the end-point population of uL16^mut^. The top 150 genes associated with impaired growth of uL16^mut^ expressing cells were then selected for further analysis.

We next performed a genome-wide fluorescence-based CRISPRi screen to identify factors mediating the degradation of uL16^mut^-RFP. We engineered a K562 cell line that constitutively expressed either the uL16^mut^-RFP or uL16^wt^-RFP reporter along with the dCas9-KRAB CRISPRi effector. Notably, the uL16^mut^-RFP cell exhibited reduced RFP fluorescence compared to the control uL16^wt^-RFP cell line (Fig. 2E), despite similar mRNA levels of the reporters (SFig. 8A). This observation is consistent with active degradation of uL16^mut^ as described in Fig. 1. We postulated that depletion of RASP factors will interfere with the ability of the cells to detect faulty biogenesis intermediates, resulting in higher RFP fluorescence. To identify such quality control factors, we transduced the uL16^mut^-RFP reporter cell line with a whole genome dual sgRNA library^68^. Subsequently, cells with high and low RFP signal were sorted via FACS (SFig. 7B) and the genes that were depleted in those cells were identified by deep sequencing the sgRNAs they expressed (Fig. 2F).

We compared the top 150 hits from both screens (SFig. 9) and identified five genes that were common in both screens: *INTS2*, *INTS3*, *S100A14*, *PRPF40A*, and *ZNF574*. *INTS2* and *INTS3* are components of the integrator complex^69^, which regulates transcription and RNA processing of various non-polyadenylated RNAs. Although the complex may be involved in the clearance of defective ribosome biogenesis intermediates, its perturbation leads to global gene expression changes^70^ and therefore many indirect phenotypes, complicating data interpretation. *S100A14,* a member of the S100 subfamily consisting of small proteins containing a Ca^2+^ binding motif ^71^, was not pursued further due to its tissue-specific cellular role and expression, as well as its plasma membrane subcellular localization^72^. The yeast homolog of *PRPF40A*, *PRP40*, is an essential splicing factor^73^, therefore, perturbations of *PRPF40A* are expected to result in global changes in gene expression, complicating studies that focus specifically on ribosome biogenesis and quality control.

We focused our attention on the poorly characterized gene *ZNF574*. Although the role of ZNF574 in ribosome biogenesis has not been formally tested, ZNF574 is predicted to have a ribosome-related role based on several high-throughput studies. In an imaging CRISPR screen, ZNF574 clustered with genes having ribosome biogenesis phenotype^74^, and several mass spectrometry studies detected physical interaction between ZNF574 and various ribosomal proteins^75–77^. Essentiality profiles derived from pan-cancer cell line CRISPR perturbations^75,76^ have revealed the uL16 chaperone protein *AAMP*^77–79^, and late acting biogenesis factor *ZNF622* as co-dependencies of *ZNF574*, as well as uL16 expression as a predictor of *ZNF574* gene essentiality. ZNF574 contains Cys2-His2 (C2H2) zinc finger binding domains that are known to interact with both DNA and RNA^80^, providing a potential mechanism for recognizing and engaging defective ribosomes. ZNF574 localizes in the nucleoplasm and cellular speckles, similar to late ribosome biogenesis factors like eIF6 and NMD3^81,82^. Finally, this protein is ubiquitously expressed^83^, and since ribosome biogenesis is a fundamental cellular process, its dedicated quality control factors are likely to be expressed in all tissues and organs.

We confirmed that depletion of ZNF574 leads to reduced cell proliferation in K562 cells expressing uL16^mut^ compared to uL16^wt^ (Fig. 2D). Additionally, we directly measured the impact of ZNF574 knockdown on uL16-RFP levels using flow cytometry (Fig. 2G). Depletion of ZNF574 (SFig 8B) preferentially stabilized uL16^mut^-RFP levels without impacting the reporter’s mRNA levels (SFig. 8C). Importantly, knocking down ZNF574 had no effect on global ribosome biogenesis (SFig. 10), confirming that ZNF574 is a novel component of RASP and not a general biogenesis factor.

### ZNF574 binds faulty pre-ribosomes and mediates their degradation

To determine whether ZNF574 binds ribosomes, we expressed GFP-tagged ZNF574 in K562 cells and performed polysome profiling, monitoring the distribution of all ribosomes (A_260_ signal) and ZNF574 (GFP signal). GFP-ZNF574 co-migrates predominantly with 60S subunits, and to a lesser extent with 40S subunits and 80S monosomes (Fig. 3A). Immunoprecipitation of GFP-tagged ZNF574 followed by mass spectrometry analysis of interacting proteins also revealed strong enrichment for ribosomal proteins from the large subunit (SFig. 11). We investigated whether ZNF574 interacts with all maturing ribosomes or preferentially binds defective ones. We co-expressed GFP-tagged ZNF574 and FLAG-tagged uL16^wt^ or uL16^mut^ in HEK293T cells. We immunoprecipitated uL16-FLAG and detected ZNF574 association via western blotting. Indeed, we detected a stronger interaction between ZNF574 and uL16^mut^ compared to uL16^wt^ (Fig. 3B), suggesting that ZNF574 targets defective large subunits.

**Fig. 3.**
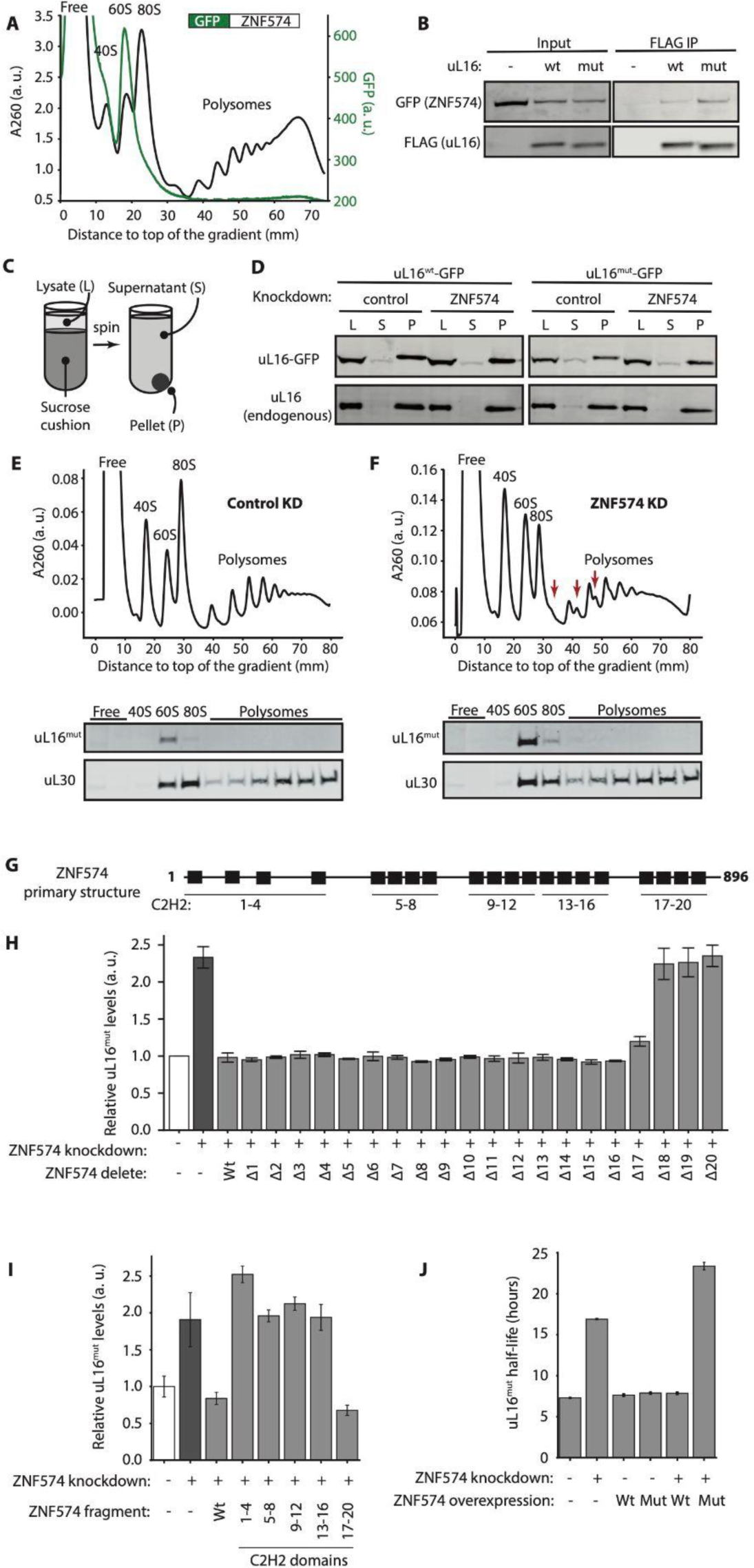
ZNF574 is a quality control factor targeting defective ribosomes. **A.** Sucrose gradient from K562 cells expressing GFP-tagged ZNF574. The total RNA absorbance at 260 nm (black line) and GFP signal (green line) were measured along the gradient. **B.** FLAG-tagged wild type or mutant uL16 was immunoprecipitated and the association with GFP-tagged ZNF574 was detected via western blot. **C.** Diagram of assay separating ribosome-associated from ribosome-free ribosomal proteins. **D.** Cell lysate (L) from cells expressing wild type or mutant uL16 was pelleted through a cushion and the abundance of uL16 in the ribosome pellet (P) versus the ribosome-free supernatant (S) was evaluated via western blot. **E,F.** Polysome gradients from K562 CRISPRi cells expressing FLAG-tagged uL16^mut^ and control sgRNA or sgRNA targeting ZNF574. Fractions were collected along the gradients and the distribution of uL16^mut^ was evaluated via western blot. uL30 was used as a loading control. **G.** Schematic of ZNF574 domain architecture showing the distribution of 20 zinc finger domains (C2H2). **H.** ZNF574 was knocked down in K562 CRISPRi cells expressing RFP-tagged uL16^mut^. Wild type or C2H2 delete ZNF574 was expressed in these cells and the uL16^mut^-RPF levels were measured via flow cytometry (mean ± SD, N=3). **I.** ZNF574 was knocked down in K562 CRISPRi cells expressing RFP-tagged uL16^mut^. Full length (Wt) ZNF574 or fragments of the protein containing the listed C2H2 domains were expressed in these cells and the uL16^mut^-RPF levels were measured via flow cytometry (mean ± SD, N=3). **J.** uL16^mut^-GFP was expressed under doxycycline inducible promoter and a pulse-chase experiment similar to Fig. 1E was performed to measure its half-life. The experiment was performed in control and ZNF574 knockdown cells that overexpress wild type ZNF574 (Wt) or ZNF574 mutant (Mut) lacking the last four C2H2 domains (mean ± SD, N=3).

Given uL16’s ability to exchange off the ribosome^84,85^, we determined whether ZNF574 mediates the release of uL16^mut^ from defective 60S subunit as a part of quality control. We took advantage of our doxycycline inducible system and expressed GFP-tagged wild type or mutant uL16 in K562 CRISPRi cells. We washed away the drug after 24 hours and started the experiment two hours after drug removal to allow full incorporation of GFP-tagged uL16 into ribosomes. To prevent rapid degradation of free ribosomal proteins^62–64^, we treated the cells with the proteasomal inhibitor Bortezomib^61^. Subsequently, we separated ribosomal proteins that are incorporated into ribosomes versus ribosome-free ribosomal proteins via ultracentrifugation (Fig. 3C). If ZNF574 extracts uL16^mut^ from the defective ribosome, we would expect an accumulation of uL16^mut^ in the ribosome-free fraction in wild type, but not ZNF574 depleted cells. Alternatively, if ZNF574 facilitates the degradation of the free uL16^mut^, we would expect an accumulation of uL16^mut^ in the ribosome-free fraction upon ZNF574 knockdown. However, we did not observe accumulation of ribosome-free uL16 under any of the tested conditions (Fig. 3D), suggesting that ZNF574 targets the entire defective 60S.

To further investigate whether ZNF574 targets the whole defective large subunit or only the free ribosomal proteins, we performed polysome profiling assays (Fig. 3E, 3F, SFig. 12). We hypothesized that if ZNF574 targets free ribosomal proteins, then knocking down ZNF574 would lead to accumulation of uL16^mut^ in the ribosome-free portion of the gradient. Conversely, if ZNF574 facilitates the degradation of the whole defective 60S, depletion of ZNF574 would lead to uL16^mut^ accumulation in the 60S fraction. Indeed, upon ZNF574 knockdown, we detected accumulation of uL16^mut^ in the 60S fraction, and no signal in the ribosome free fraction (Fig. 3F). Together, our data show that ZNF574 functions as a quality control factor that targets defective large subunits and not free ribosomal proteins.

### The last four C2H2 domains of ZNF574 are necessary and sufficient for uL16^mut^ degradation

We next aimed to identify the regions within the ZNF574 protein involved in the recognition of faulty ribosomes containing uL6^mut^. Unfortunately, our efforts were hindered by a lack of structural information for ZNF574. Even structure prediction algorithms, such as AlphaFold^86,87^, were unable to provide a confident tertiary structure for the full-length ZNF574 (SFig. 13). ZNF574 is an approximately 99 kDa zinc finger protein^88^ containing 20 C2H2 domains (Fig. 3G) that can interact with DNA, RNA, or proteins^80^. To systematically investigate the role of each C2H2 domain, we performed deletion studies by removing individual C2H2 domains and testing the ability of the obtained mutants to degrade uL16^mut^ (Fig. 3H, SFig. 14). We knocked down the endogenous ZNF574 in K562 CRISPRi cells expressing uL16^mut^-RFP, which led to stabilization of the defective ribosomes (Fig. 3H). We then introduced either a wild type ZNF574 or a C2H2-deleted variant and measured uL16^mut^-RFP levels via flow cytometry to determine whether these constructs would rescue uL16^mut^-RFP degradation. Our findings revealed that the last four C2H2 domains of ZNF574 are critical for uL16^mut^ degradation and deleting any one of them interferes with ZNF574-mediated quality control.

Next, we tested whether a specific region of the protein is sufficient for uL16^mut^ degradation. We introduced either a GFP-tagged full length ZNF574 or a fragment of the protein containing a group of four C2H2 domains in ZNF574 depleted K562 cells expressing uL16^mut^-RFP (Fig. 3I). Surprisingly, introducing the last four C2H2 domains (C2H2 17-20) restored uL16^mut^-RFP degradation to the same extent as the full-length protein. Taken together, these data demonstrate that the C-terminus of ZNF574 is both necessary and sufficient for uL16^mut^ degradation.

To gain insight into ZNF574-mediated clearance of uL16^mut^, we performed a pulse-chase experiment where we expressed uL16^mut^-GFP under a doxycycline inducible promoter^48^ in control cells or ZNF574 knockdown cells. Upon inducing uL16^mut^ expression for 24 hours, we washed away the doxycycline, and measured the levels of uL16^mut^-GFP via flow cytometry (Fig. 3J). Our results show that knockdown of ZNF574 dramatically increased the half-life of uL16^mut^ from approximately 7 hours to approximately 17 hours, confirming the pivotal role of ZNF574 in uL16^mut^ degradation. We then overexpressed GFP-tagged wild type ZNF574 or mutant ZNF574 lacking the last four C2H2 domains (ZNF574^mut^) and measured uL16^mut^ half-life. In control cells, overexpression of ZNF574 did not accelerate uL16^mut^ degradation, suggesting that despite the low endogenous levels, ZNF574 is not rate-limiting for faulty ribosome clearance. Furthermore, overexpression of ZNF574^mut^ in cells that express wild type endogenous ZNF574 also did not impact uL16^mut^ degradation rates. Finally, we performed rescue experiments where we introduced wild type or mutant ZNF574 in ZNF574 knockdown cells. Overexpression of wild type, but not mutant ZNF574 restored uL16^mut^ degradation in these cells (Fig. 3J).

### Loss of ZNF574 gradually leads to shutdown of 60S biogenesis

We observed that loss of ZNF574 in cells expressing uL16^mut^ led to abnormal polysomes, as indicated by shoulders on the monosome, disome, and trisome peaks (Fig. 3F, red arrows). These secondary peaks, known as half-mers, consist of polysomes containing 80S ribosomes and a single preinitiation complex comprised of a 40S subunit plus initiation factors^89^. Since ribosome initiation is a rapid process^90^, half-mers are typically observed when production of large subunits is impaired or there is a defect in subunit joining^91^. We hypothesized that loss of ZNF574 in combination with higher levels of ribosome biogenesis defects due to the presence of uL16^mut^ ultimately leads to global perturbations in ribosome biogenesis. Indeed, trapped biogenesis intermediates contain ribosome biogenesis factors, such as eIF6 and NMD3 (Fig 1C, D). Since these factors are expressed at low levels^83^ and serve numerous ribosomes, their reduced availability due to entrapment on defective pre-ribosomes can inhibit ribosome production. To validate this hypothesis, we compared eIF6’s subcellular localization in U2OS CRISPRi cells that express wild type or mutant uL16 as well as sgRNA targeting ZNF574 (Fig. 4A). Although eIF6 dynamically shuttles between the nucleus and the cytoplasm with large subunit biogenesis intermediates, the bulk of the protein is typically localized within the nucleus^92^. However, upon ZNF574 depletion and uL16^mut^ expression, eIF6 was predominantly localized in the cytoplasm (Fig. 4A, B). This observation confirms our hypothesis that perturbing RASP leads to accumulation of defective biogenesis intermediates still bound to biogenesis factors, effectively trapping them in the cytoplasm. The loss of these biogenesis factors from the nucleus interferes with normal ribosome assembly.

**Fig. 4.**
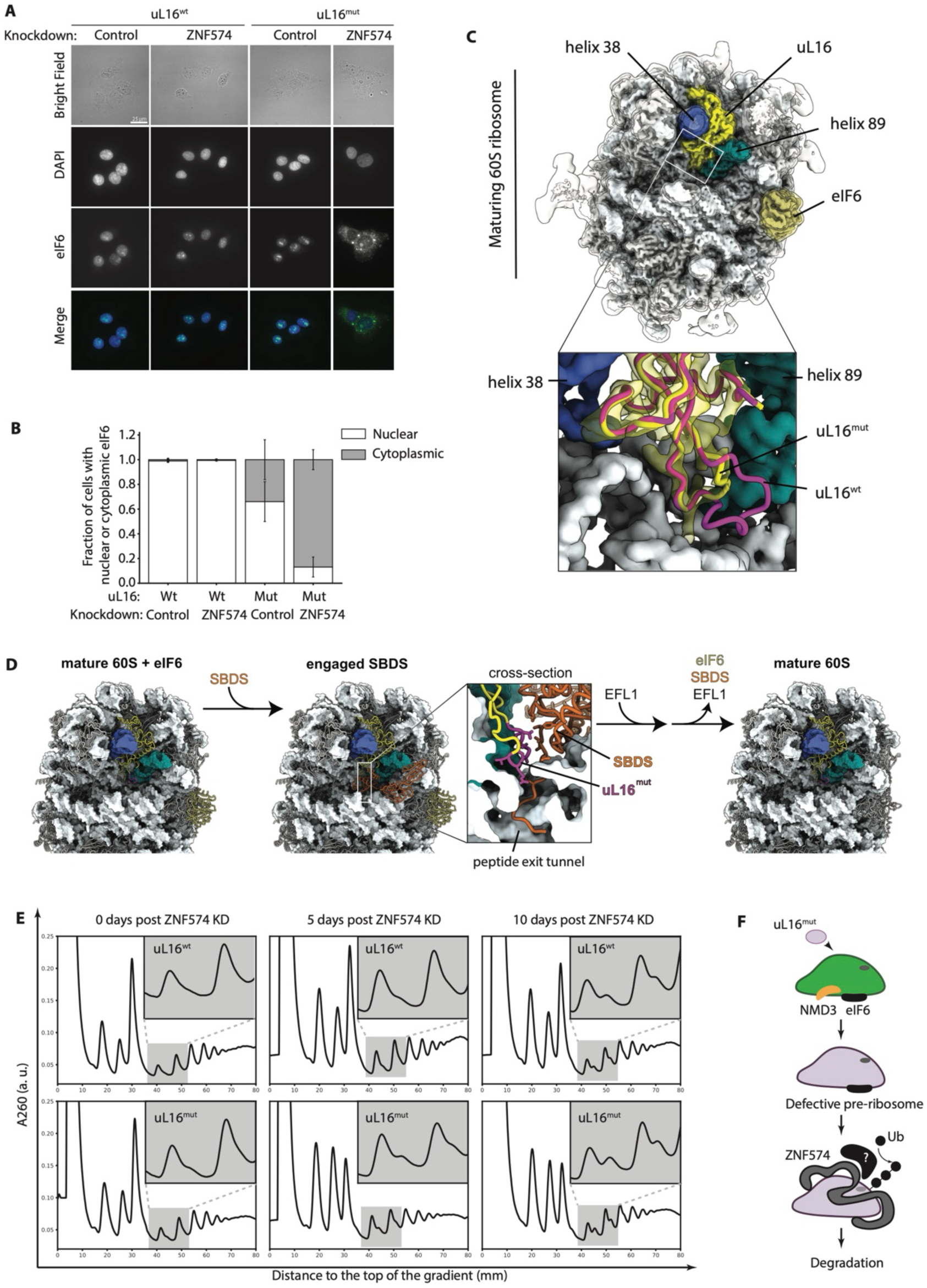
Loss of quality control leads to global defects in ribosome production. **A.** ZNF574 was knocked down in U2OS CRISPRi cells stably expressing wild type or mutant uL16 and the subcellular localization of eIF6 was examined by fluorescence microscopy. Scale bar, 25 µm. **B.** The fraction of cells with predominantly nuclear or cytoplasmic eIF6 was manually calculated under each condition (mean ± SD, N=3). Data was calculated from three biological replicates where approximately 600 cells were evaluated per condition. **C.** Cryo-EM reconstruction of mature human pre-60S with incorporated uL16^mut^ filtered to a resolution of 8 Å. Box: close-up of uL16^mut^ density and comparison of the built model with uL16^wt^ from PDB 4ug0. **D.** Simplified scheme for the last step of 60S maturation, which removes eIF6 from the mature large subunit through the action of SBDS and EFL1. Box: cross-section of the ribosome with bound SBDS to show the interaction of SBDS with the P-loop of uL16. Highlighted in magenta are those residues that were deleted in uL16^mut^. The displayed models were prepared using the refined structure of human 60S and the deposited coordinates for SBDS bound to 60S subunits from PDB 5an9. **E.** ZNF574 was knocked down in K562 CRISPRi cells expressing wild type or mutant uL16 and samples were harvested and analyzed on polysome gradients 0, 5, or 10 days post knockdown. Inserts on the upper right of each graph represent a zoom-in of the disomes and trisomes (shaded rectangle), where half-mers first appear. **F.** Model for ZNF574-mediated ribosome quality control.

We confirmed that the association of cytoplasmic eIF6 is bound to the defective large subunits by comparing its migration on polysome gradients upon ZNF574 depletion and uL16^mut^ expression (SFig. 15). We then purified 60S subunits from K562 CRISPRi cells expressing uL16^mut^ and sgRNA targeting ZNF574 and investigated them using cryo-electron microscopy. Indeed, all 60S particles containing uL16^mut^ were bound to eIF6. Our reconstruction of 60S^mut^ bound to eIF6 achieved a resolution of 2.5 Å, allowing us to determine the structure of uL16^mut^ by direct inspection of the cryo-EM map density (Fig. 4C).

The structure of uL16^mut^ in context of the 60S subunit suggests a mechanism for the retention of eIF6 on 60S^mut^. Eviction of the antiassociation factor eIF6 from 60S subunits is considered the last step of 60S maturation and requires the coordinated action of the proteins SBDS and EFL1. Cryo-EM studies of human SBDS bound to *Dictyostelium discoideum* pre-60S revealed contacts of SBDS domain I with the P-site loop of uL16^38^. It has been postulated that some mutations of uL16 that destabilize or disrupt its P-site loop may prevent the release of eIF6 by perturbing the binding of SBDS to pre-60S^37,38^. This concept is in good agreement with our observation that eIF6 is bound to all our pre-60S particles carrying the uL16:102–111 deletion, which would abolish interactions of the P-site loop of uL16 with SBDS (Fig. 4D). The inability to evict eIF6 also explains the accumulation of eIF6 in the cytoplasm of cells with ZNF574 knockdown expressing uL16^mut^ (Fig. 4A,B).

We hypothesized that the loss of ZNF574 may also impact ribosome biogenesis in wild type cells lacking uL16^mut^ expression. However, we expected that the sequestration of biogenesis factors in the cytoplasm and appearance of half-mers would occur more gradually, since it depends on accumulation of stochastic errors during ribosome assembly. To investigate this hypothesis, we knocked down ZNF574 in cells expressing either wild type or mutant uL16 and monitored the appearance of half-mers over time via polysome gradients (Fig. 4E). Cells expressing uL16^mut^ accumulated half-mers five days after ZNF574 depletion. However, cells expressing uL16^wt^ did not exhibit half-mers five days post ZNF574 depletion, but displayed them after ten days. This experiment shows the inherent error-prone nature of ribosome maturation, and suggests that inefficient clearance of defective biogenesis intermediates, as observed in ZNF574 knockdown, can sequester biogenesis factors, ultimately leading to inhibition of ribosome production (Fig. 4F).

## Discussion

The essential process of ribosome assembly is vulnerable to environmental perturbations and stochastic errors as a consequence of its inherent complexity. To ensure uninterrupted ribosome production and protein synthesis, cells must detect and eliminate improperly assembled biogenesis intermediates. Despite the critical role of these quality control mechanisms, the factors involved in monitoring ribosome assembly have remained elusive. In this study, we modeled stochastic errors in large subunit assembly and identified a novel factor, *ZNF574*, that targets defective large subunits for proteasomal degradation. Our findings demonstrate the essential role of *ZNF574* and the ribosome assembly surveillance pathways (RASP) it contributes to in sustaining ribosome production and cellular health.

How does ZNF574 mediate ribosome degradation? We discovered that faulty biogenesis intermediates undergo degradation via the ubiquitin-proteasome system (Fig. 1G). Although ZNF574 is not itself an E3 ligase, previous studies have demonstrated that zinc finger proteins can serve as scaffolds to recruit enzymatically active complexes^93,94^. Thus, it is possible that ZNF574 initiates the recruitment of E3 ligases and other factors, including machinery to degrade the ribosomal RNA that is part of the faulty ribosome. Our mass spectrometry studies on ZNF574 or uL16^mut^ did not yield any candidate interactors (SFig. 11, Fig. 1D), likely due to the transient nature of these interactions, which occur only upon detection of a faulty ribosome by ZNF574. Alternatively, ZNF574 can recruit other modifying enzymes and initiate a cascade of events via modifications, such as phosphorylation, SUMOylation, or UFMylation of ribosomal proteins^95,96^, or modifications on the rRNA^97^. Under such conditions, the binding of the processing factors is sequential, and may not be detected using conventional assays such as co-immunoprecipitation and mass spectrometry. Furthermore, our CRISPRi screens are data-rich and contain numerous intriguing candidates, which role in quality control and their underlying molecular mechanism require further exploration.

Is ZNF574 a quality control factor solely dedicated to uL16? The large subunit ribosome biogenesis is blocked upon incorporation of uL16^mut^ (Fig. 1B) and such trapped intermediates require ZNF574 for degradation (Fig. 2G). Surprisingly, only the last four C2H2 domains of ZNF574 are necessary and sufficient for degradation of large subunits containing uL16^mut^ (Fig. 3H, I). Intriguingly, the predicted structure of ZNF574 implies that this C-terminal region may form a structured tertiary fold (SFig. 13C), whereas most regions of ZNF574 are not predicted to form long-range interactions. We hypothesize that different C2H2 groups of ZNF574 monitor various ribosome biogenesis intermediates, as evidenced by ZNF574’s co-migration with several ribosome species on polysome gradients (Fig. 3A), and its co-immunoprecipitation with various biogenesis factors (SFig. 11). In general, zinc finger proteins represent a large but poorly characterized group of proteins that has traditionally been associated with transcription^98^. However, C2H2 domains can also facilitate binding to RNA and other proteins^80^, making zinc finger proteins versatile components of complex biological pathways, such as ribosome biogenesis (*ZNF574*, *ZNF622*) or DNA repair^93^. Therefore, ZNF574 is likely to be a representative member of a new class of C2H2 zinc finger proteins in RASP.

Given the complexity of ribosome assembly, which requires numerous transformations of the pre-ribosomes, including incorporation of ribosomal proteins, trimming, modifying, and folding of the rRNA^99^, it is unlikely that a single factor can patrol all stages of ribosome biogenesis and differentiate between on-path and defective intermediates. Instead, various checkpoints along the ribosome assembly line most likely employ dedicated quality control factors. Our study lays the experimental framework for identifying and characterizing new RASP factors by systematically modeling ribosome biogenesis defects along the assembly pathway. By introducing targeted mutations in different ribosomal proteins, we can impede specific steps of biogenesis and visualize and purify the resulting faulty pre-ribosomes. Importantly, integrating such experimental setup with unbiased whole genome screens, such as the CRISPRi screens described in Fig. 2, allows us to pinpoint the specific quality control factors responsible for monitoring each step of ribosome assembly and maturation. Through systematic investigation of critical stages in ribosome formation, we can gain a comprehensive understanding of RASP and its role in maintaining uninterrupted ribosome production.

Furthermore, our findings demonstrate the critical role of RASP for cellular and organismal health. Loss of ZNF574 decreased ribosome production via sequestration of assembly factors on the defective ribosomes (Fig. 4), highlighting the importance of RASP. In patients, ribosome biogenesis defects lead to a group of disorders called ribosomopathies, characterized by severe anemias, cancer predisposition, skeletal abnormalities, and growth retardation^18^. While ribosomopathies have traditionally been associated with mutations in ribosomal proteins or biogenesis factors^20^, our work predicts that mutations in RASP factors, including *ZNF574*, may lead to ribosomopathy-like phenotypes in patients. Therefore, studying this new surveillance pathway will further our understanding of fundamental cell biology, which can serve as the foundation for improving patient diagnostics, and inspire novel therapeutic interventions.

## Acknowledgements

We thank A. Johnson, R. Green, J. Black, M. Catapovic, N. Sinha and members of the Green Lab for their scientific input and helpful discussions.

## Author Contributions

J.A and K.K. conceived the study. J.A., H.S., C.J, Y.W., T.G., T.H, and K.K. performed the experiments described in Fig. 1-4. S.-L.X., A.V. R. and T.G. performed mass spectrometry data acquisition and analysis. A.B and N.B. performed cryo-EM data acquisition and analysis. K.K. wrote the manuscript with input from all authors.

---

**SFig. 1.**
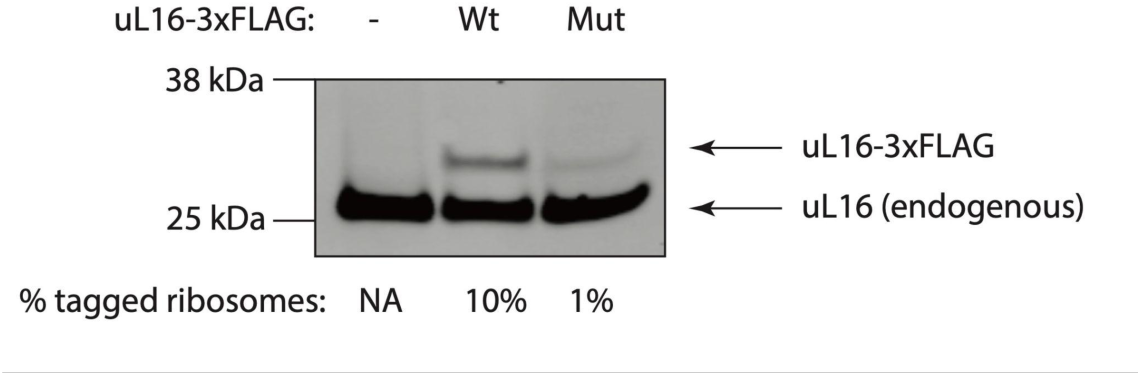
Expression levels of FLAG-tagged uL16. K562 cells were transduced with FLAG-tagged wild type or mutant uL16. The levels of the ectopically expressed and endogenous uL16 were compared via western blot with uL16 antibody. The fraction of tagged uL16 was estimated by measuring the intensity of the uL16-3xFLAG band versus the uL16 endogenous band using the LiCOR Image Studio Software.

**SFig. 2.**
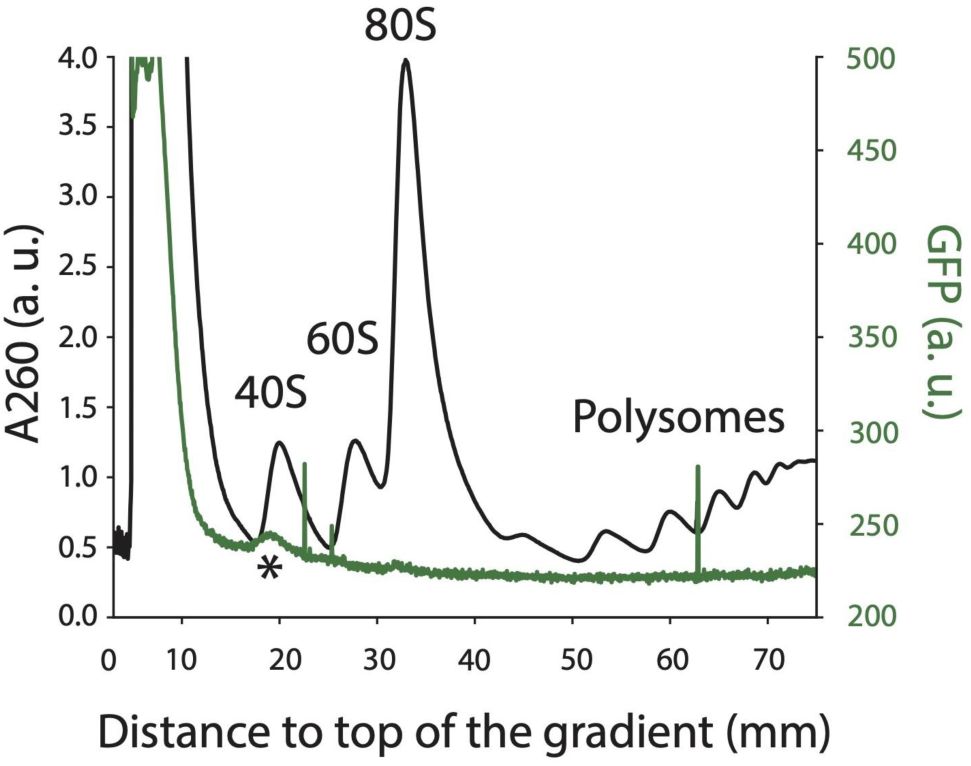
Polysome gradients from K562 cells. Sucrose gradient from untransduced K562 cells. The total RNA absorbance at A260 nm (black line) and GFP signal (green line) were measured along the gradient. The peak (*) around the 40S subunit represents auto-fluorescent signal.

**SFig. 3.**
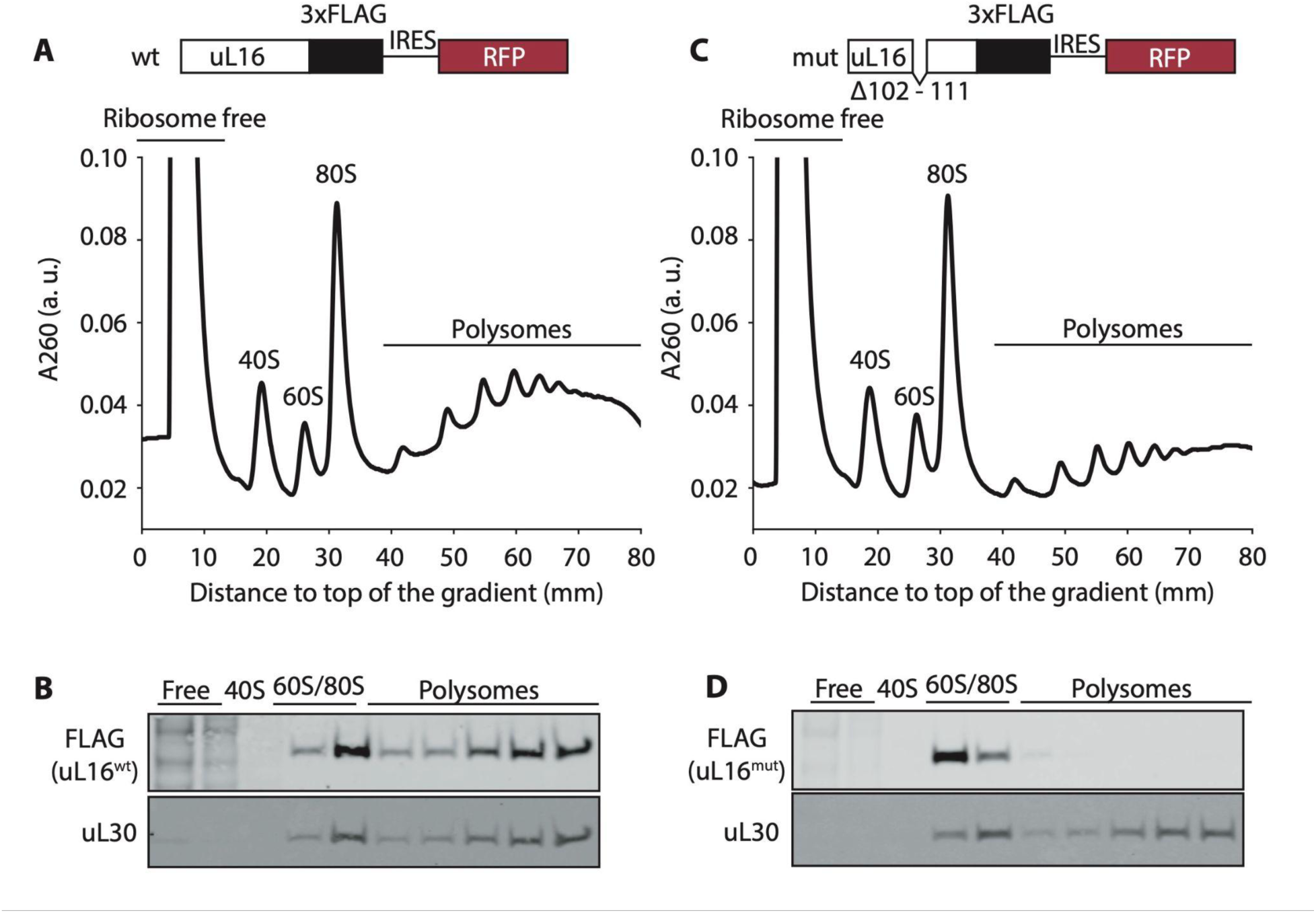
Characterization of FLAG-tagged uL16. **A, C.** Top: Diagram of wild type and mutant uL16 constructs transduced in K562 cells. Bottom: Sucrose gradients from K562 cells transduced with the indicated constructs. **B, D.** Western blots from polysome fractions corresponding to ribosome-free (free) cell lysate, ribosomal subunits (40S, 60S), monosomes (80S) and polysomes. Note that the lanes labeled 60S/80S contain fractions that capture both the 60S and 80S peaks and cannot be clearly assigned to a unique ribosome population. The distribution of uL16 wild type or mutant was determined by probing with a FLAG antibody. uL30 was used as a loading control.

**SFig. 4.**
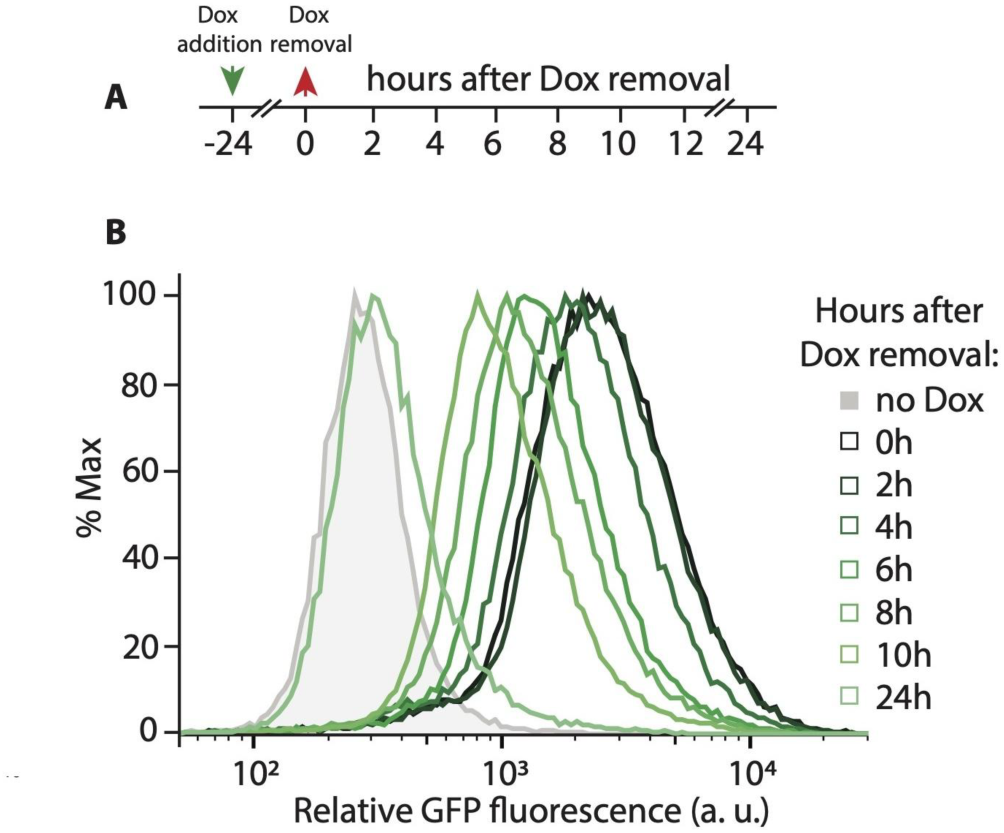
uL16-GFP pulse-chase. **A.** Diagram of experimental time-course. Expression of uL16^mut^-GFP was induced for 24h and flow cytometry measurements were conducted every 2 hours after washing out the drug for the first 10h and then again after a total of 24h. **B.** Representative flow-cytometry data from K562 cells expressing uL16^mut^-GFP under a doxycycline-inducible promoter.

**SFig. 5.**
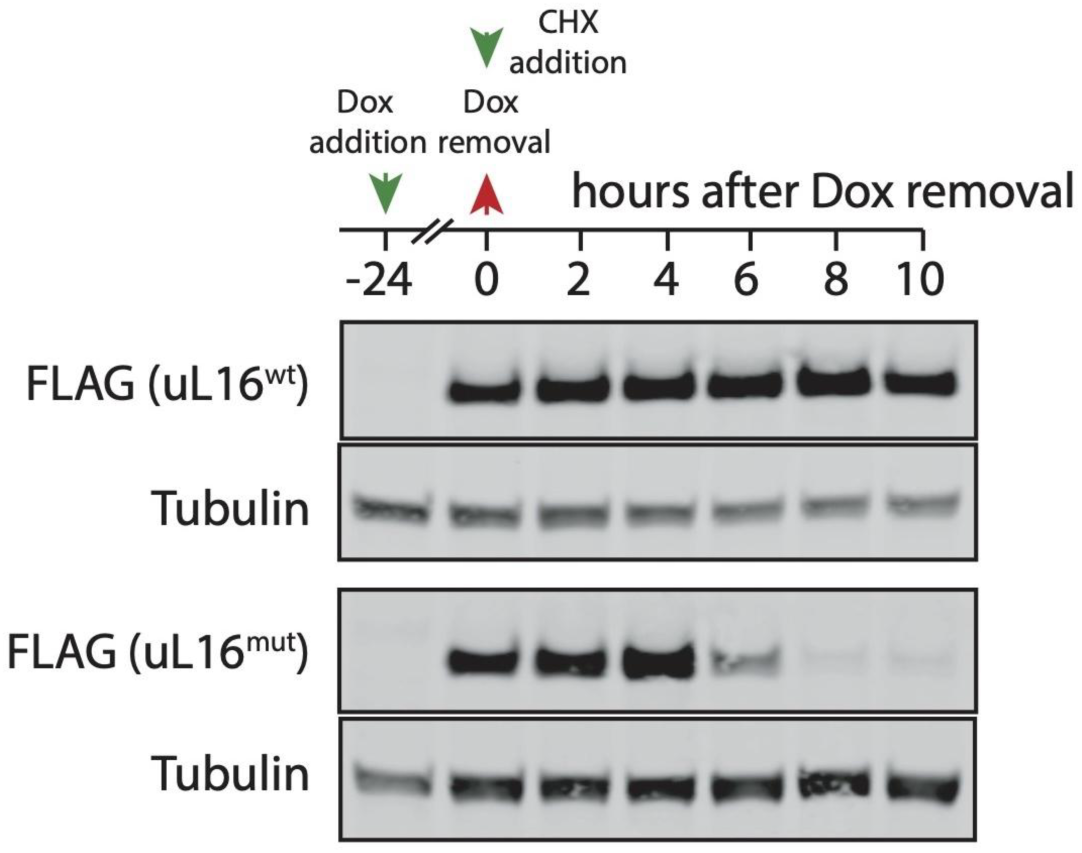
uL16^mut^ degradation does not require active translation. K562 cells expressing wild type (wt) or mutant (mut) uL16-FLAG under a doxycycline inducible promoter were cultured in the presence of doxycycline (Dox) for 24h. The drug was washed away and the translation elongation inhibitor cycloheximide (CHX) was added to the media at time point 0 h (diagram above). Samples were collected for western blot analysis at the indicated time points. Tubulin was used as a loading control.

**SFig. 6.**
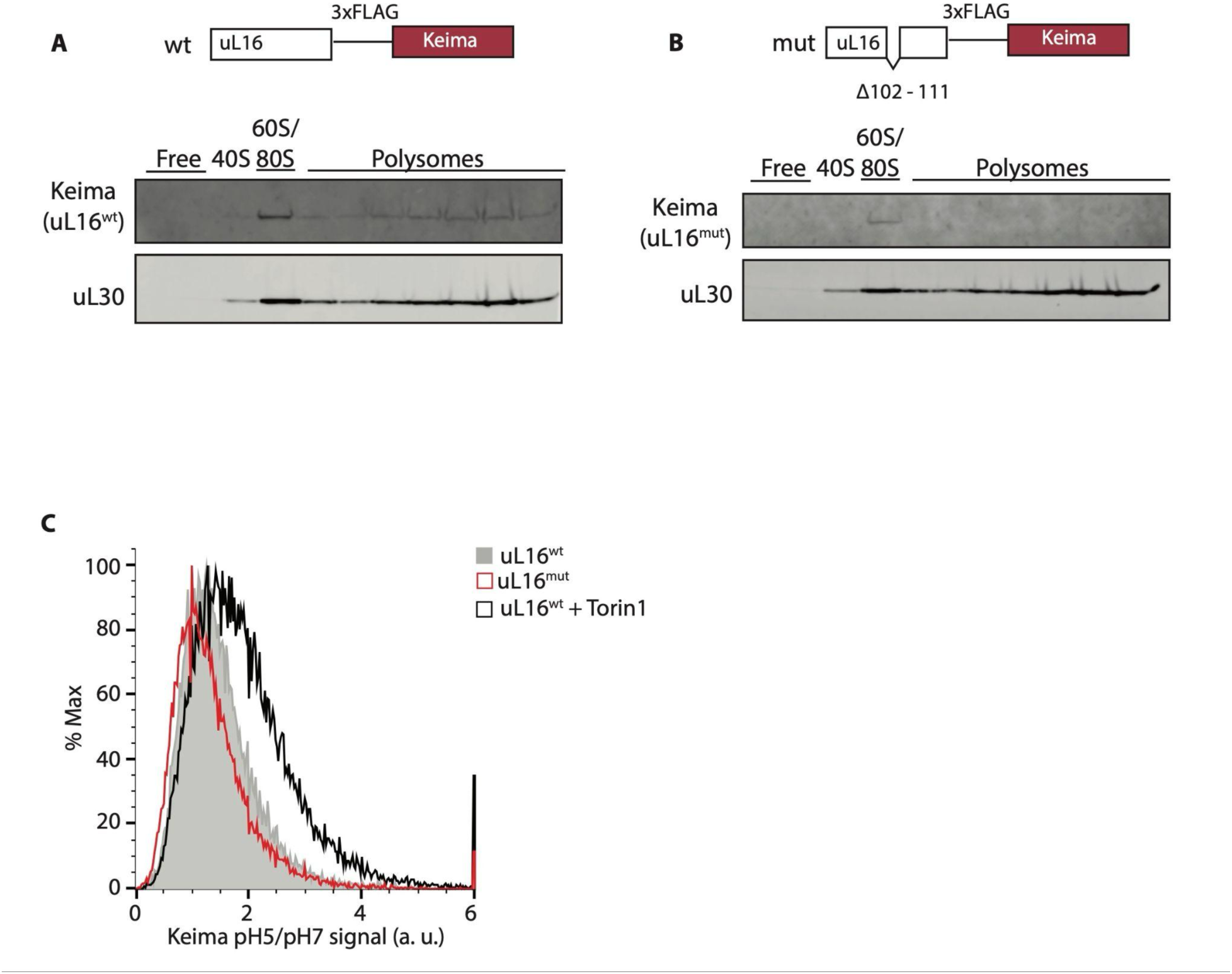
uL16^mut^ is not degraded via autophagy. **A, B.** Top: Diagrams of wild type and mutant uL16 constructs transduced in K562 cells. Bottom: Western blots from polysome fractions corresponding to ribosome-free (free) cell lysate, 40S ribosomal subunits, combined 60S and 80S fractions (60S/80S) and polysomes. The distribution of uL16 wild type or mutant was determine by probing with a Keima antibody. uL30 was used as a loading control. **C.** Keima-tagged wild type or mutant uL16 were transduced in K562 cells and their targeting to the lysosome was measured by the Keima pH5/pH7 signal. Treatment with Torin1, which induces autophagy, was used as a positive control.

**SFig. 7.**
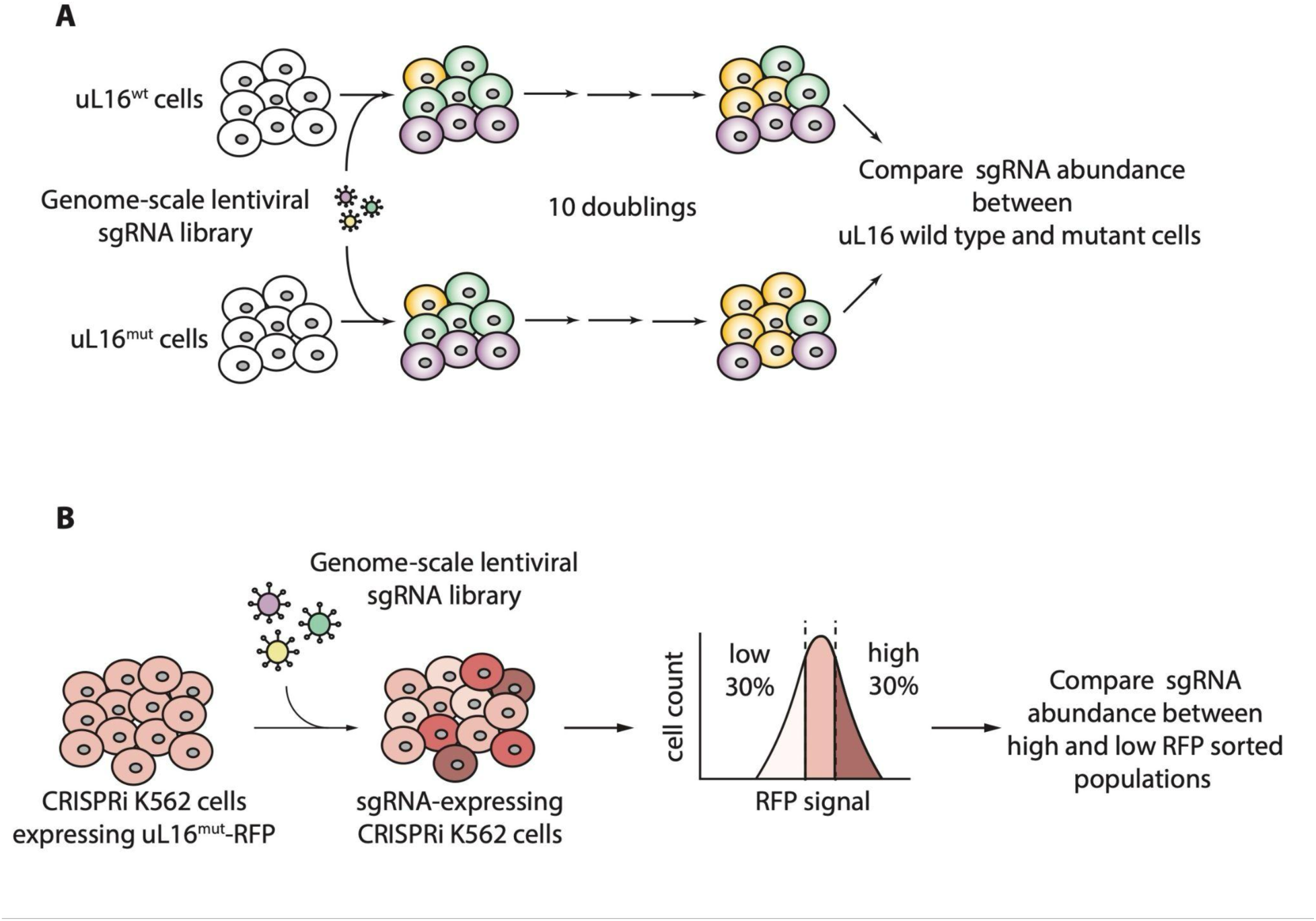
Diagram of CRISPRi screens. **A.** Growth based CRISPRi screen. **B.** FACS based CRISPRi screen.

**SFig. 8.**
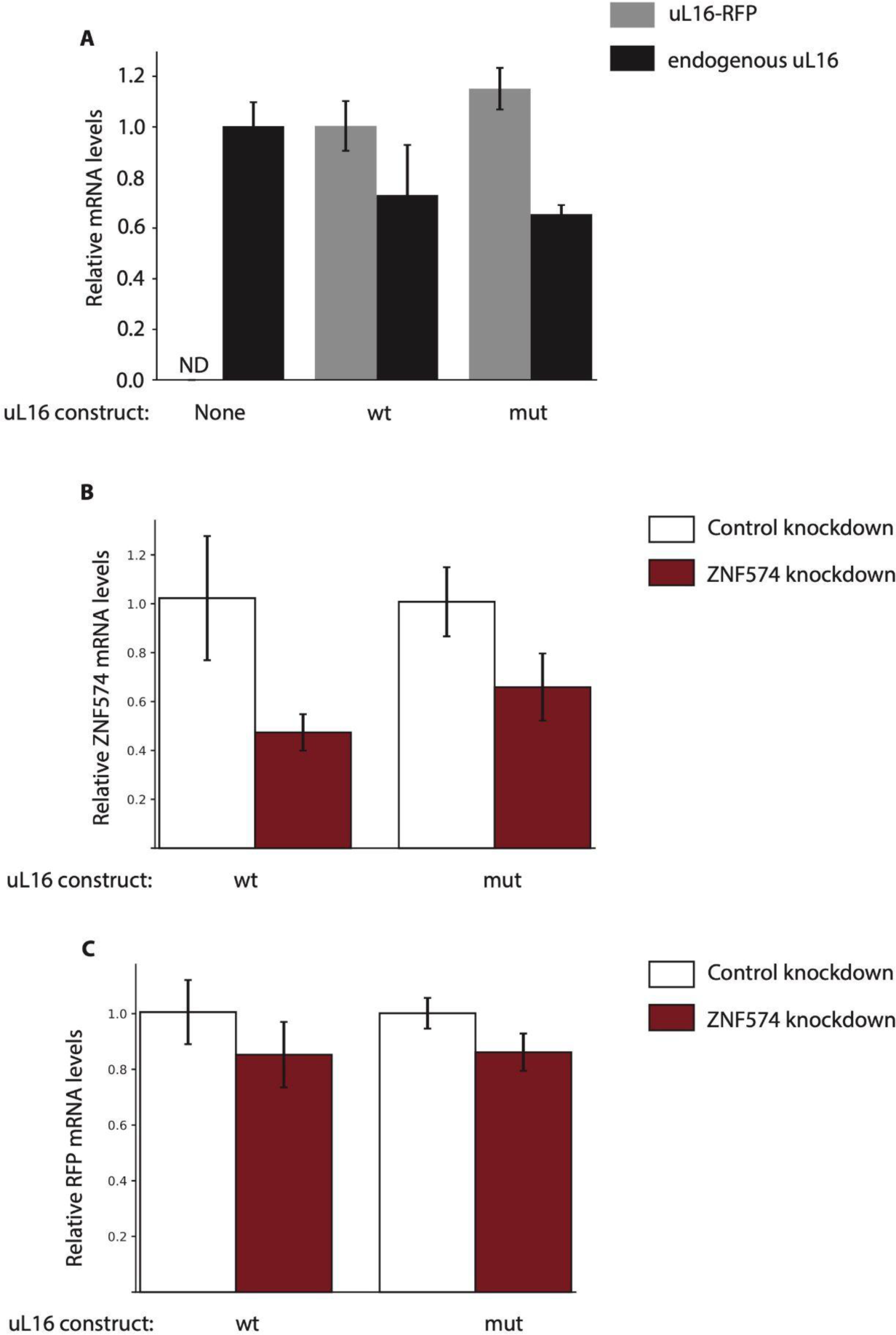
Measurement of transcript levels in wild type and knockdown K562 cells. mRNA concentrations were measured using quantitative polymerase chain reaction (qPCR) **A.** mRNA levels of endogenous and ectopic uL16 in control cells or cells transduced with wild type or mutant uL16-RFP (mean ± SD, N=3). Endogenous uL16 levels are normalized to uL16 in untransduced cells. uL16-RFP levels are normalized to uL16^wt^-RFP. **B.** mRNA levels of ZNF574 in K562 CRISPRi cells expressing a control sgRNA or sgRNA targeting ZNF574 (mean ± SD, N=3). **C.** mRNA levels of uL16-RFP reporter in K562 CRISPRi cells expressing a control sgRNA or sgRNA targeting ZNF574 (mean ± SD, N=3).

**SFig. 9.**
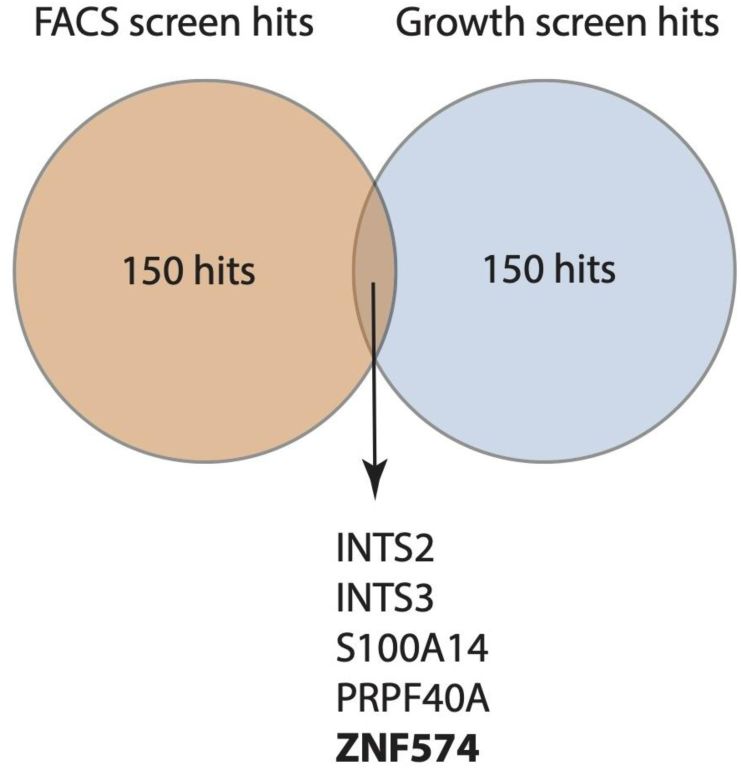
CRISPRi screens overlap. Venn diagram showing the overlap among the top 150 hits from the two CRISPRi screens described in Fig. 2.

**SFig. 10.**
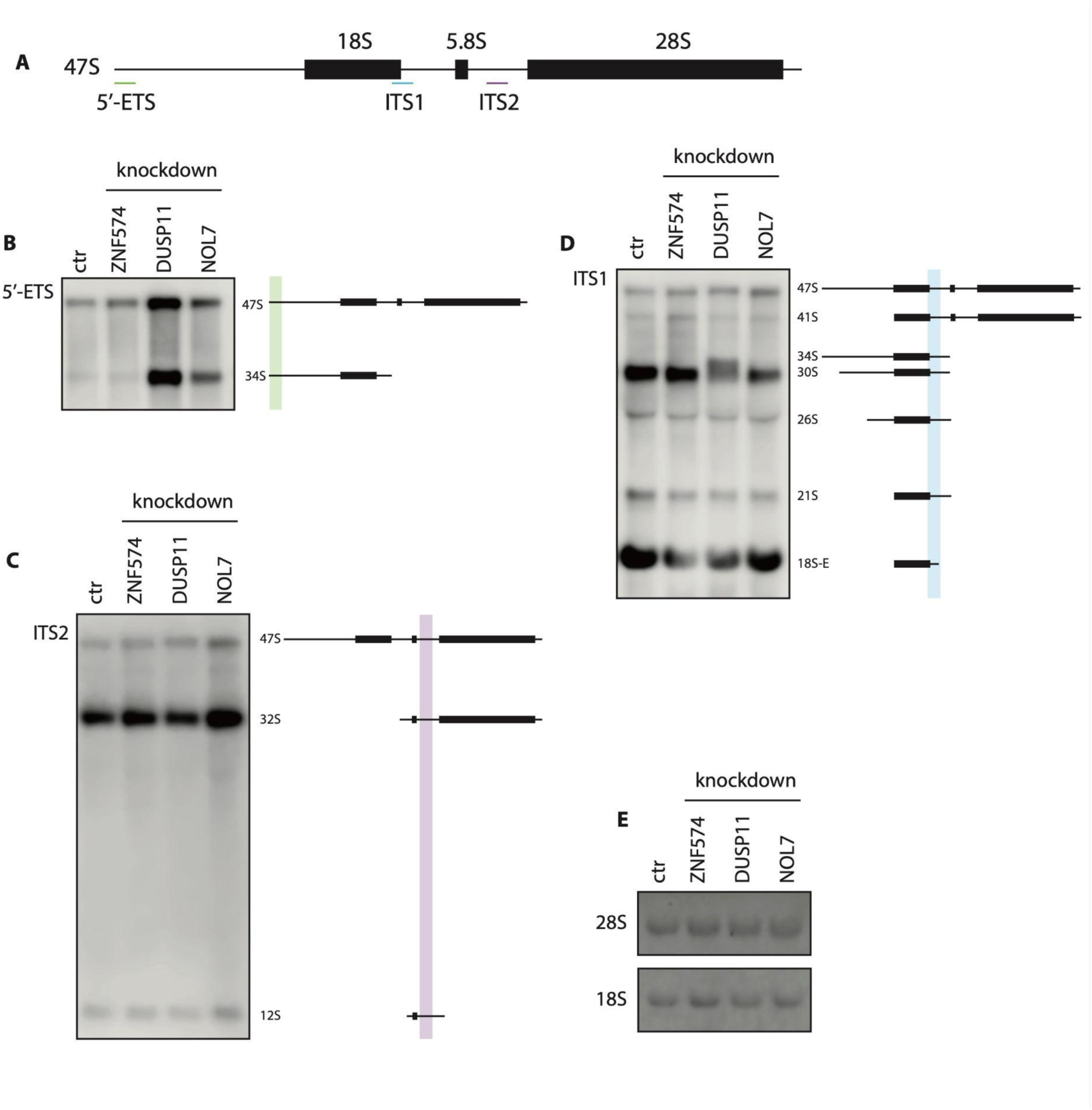
ZNF574 depletion does not perturb ribosomal RNA processing. **A.** Diagram of the 45S ribosomal RNA precursor and binding sites for the three Northern blot probes. **B-D**. Northern blot showing ribosomal RNA processing in control K562 CRISPRi cells or ZNF574, DUSP11 or NOL7 knockdown cells. Diagramed on the right are the detected processing intermediates. DUSP11 and NOL7 are proteins important for ribosomal RNA processing and are used as positive controls. **E.** Ethidium bromide staining of the 18S and 28S mature ribosomal RNA is used as a loading control.

**SFig. 11.**
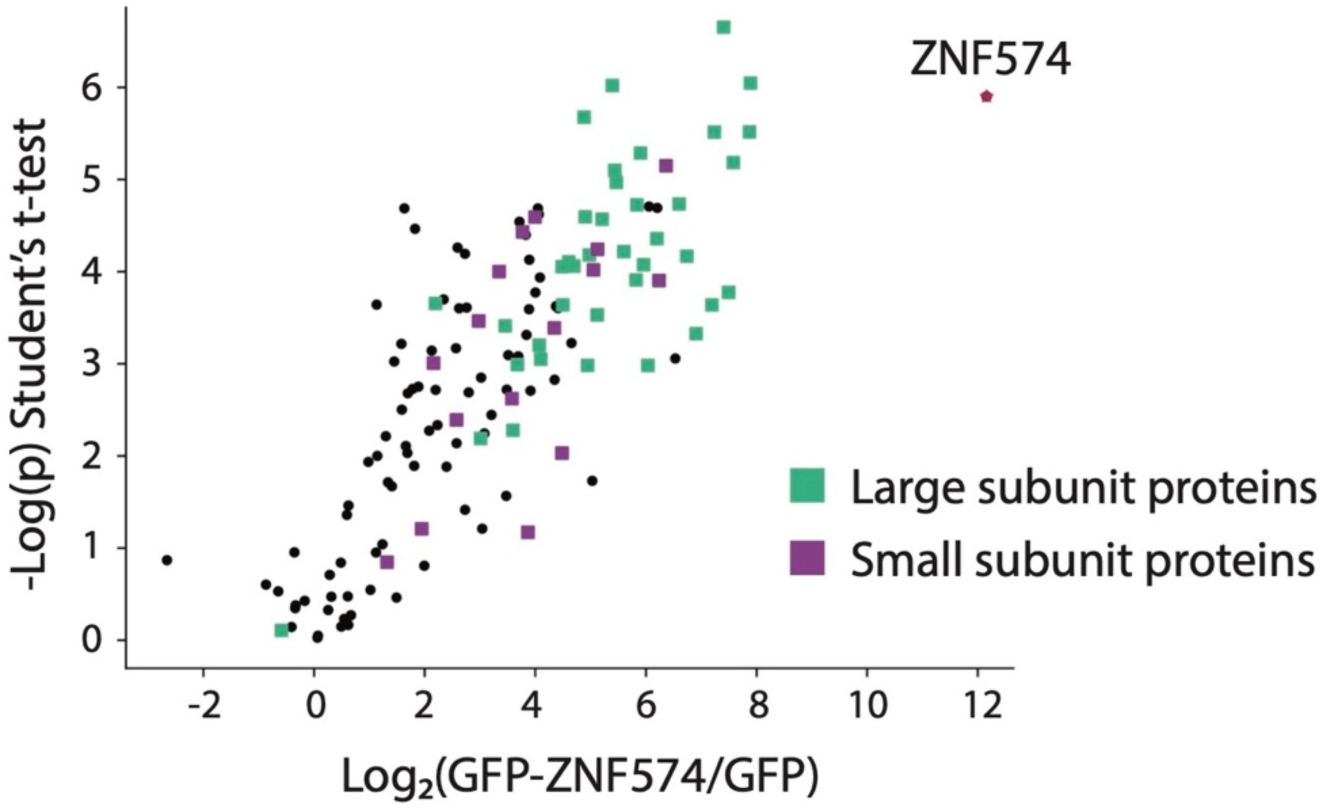
ZNF574 binds ribosomes. GFP-tagged ZNF574 or GFP were expressed in 293T cells. The proteins were immunoprecipitated and the associated proteins were identified via mass spectrometry. The graph plots the enrichment of proteins in the GFP-ZNF574 co-immunoprecipitated sample. Highlighted are ribosomal proteins from the large subunit (green square), small subunit (purple square), and ZNF574 (red star).

**SFig. 12.**
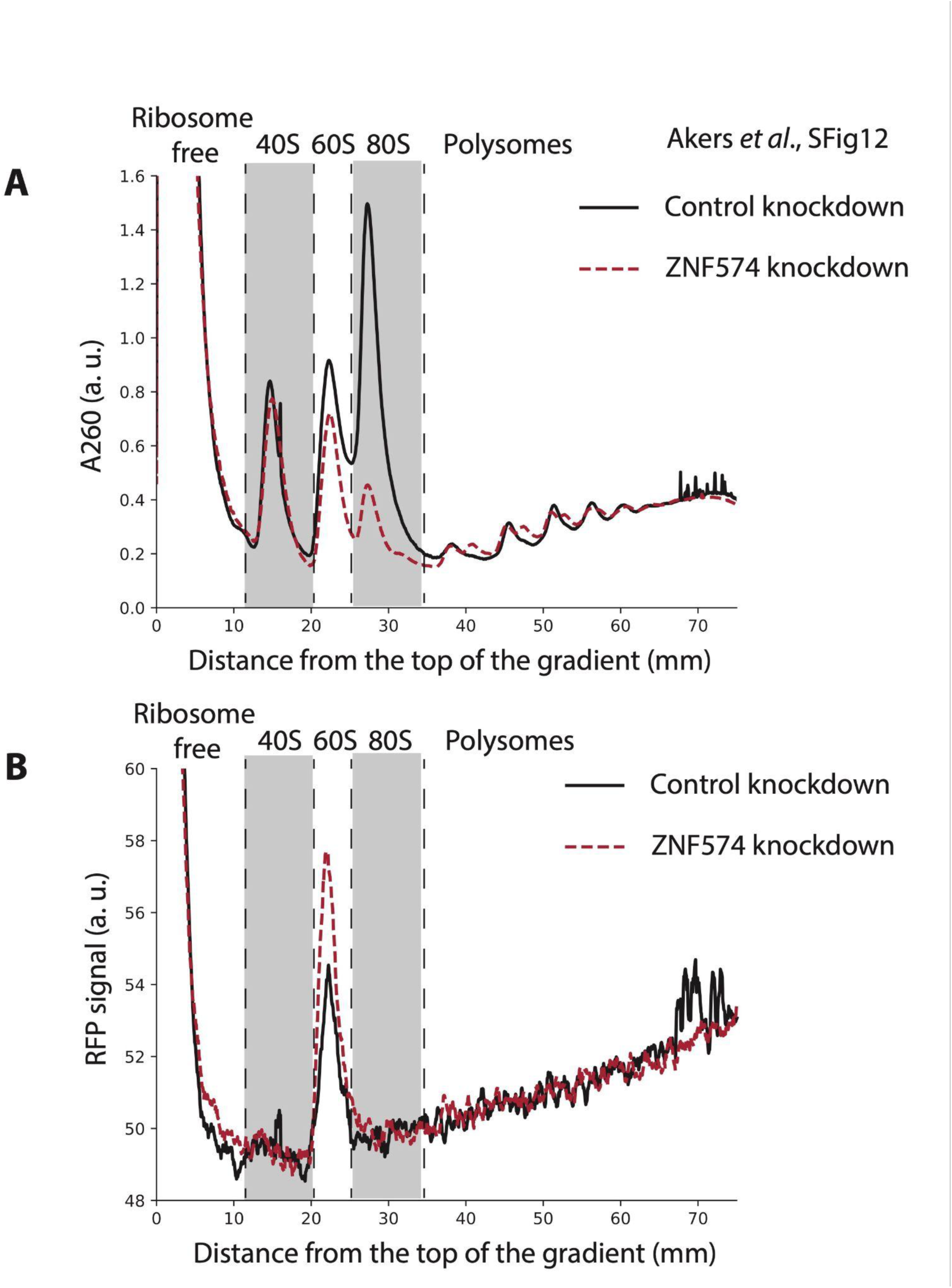
Knocking down ZNF574 stabilizes defective large subunits and not ribosome-free uL16^mut^. **A.** Polysome gradients from K562 CRISPRi cells expressing uL16^mut^-RFP and control sgRNA (black line) or sgRNA targeting ZNF574 (red line). **B.** RFP signal from the same gradients as in A. Highlighted are the ribosome free fraction and the 60S fraction. Knocking down ZNF574 increases uL16^mut^-RFP signal specifically in the 60S fraction.

**SFig. 13.**
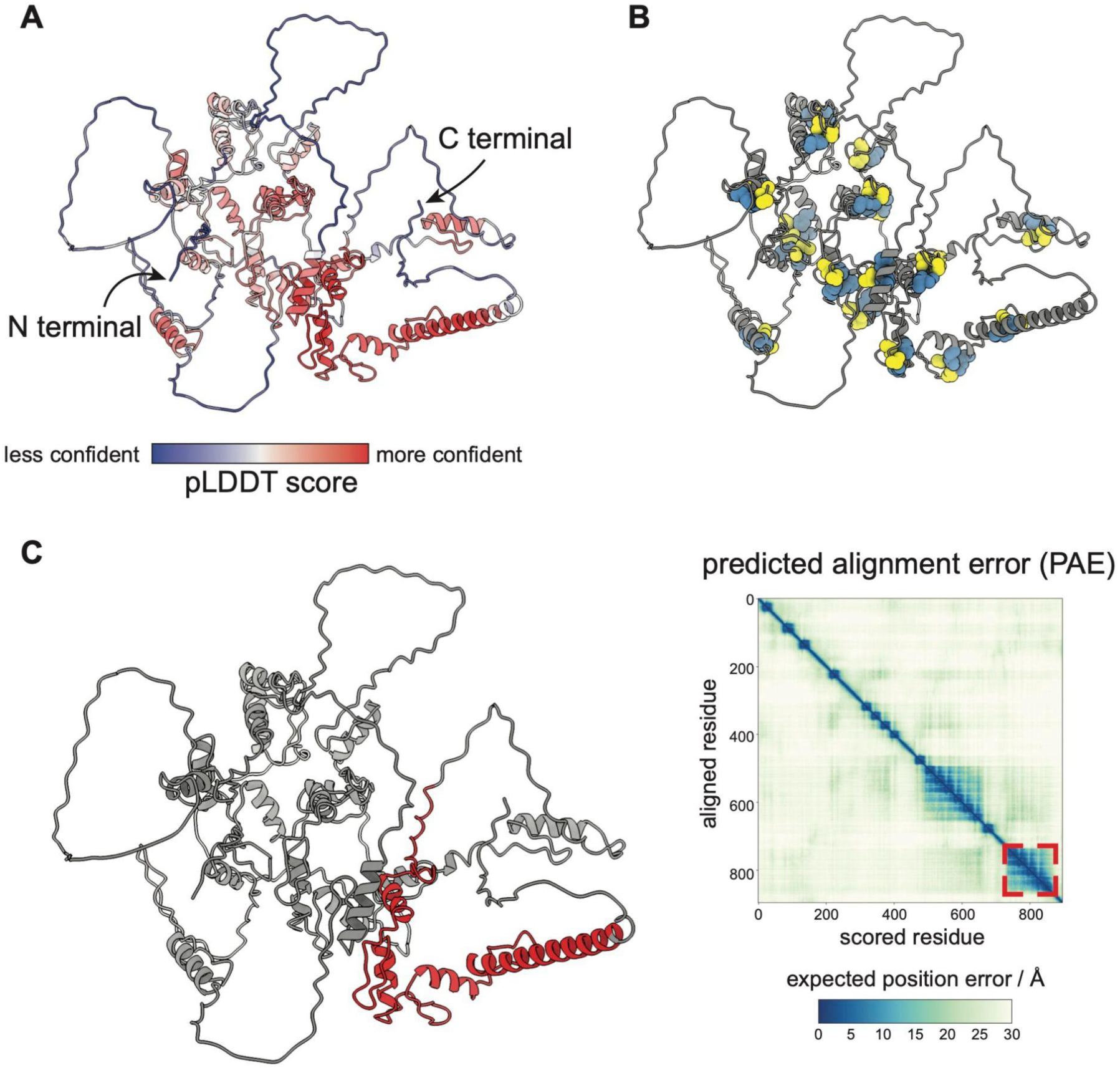
Structure model of ZNF574 as predicted by AlphaFold. **A.** Three-dimensional model of human ZNF574 as predicted by the Alphafold algorithm. The confidence is color-coded (see scale below). **B.** The same model highlighting the 20 zinc finger (C2H2) motifs shown as spheres (yellow: cysteine residues, blue: histidine residues). **C.** The analysis of the predicted alignment error (PAE) identifies a putatively folded C-terminal domain comprising the last four zinc fingers.

**SFig. 14.**
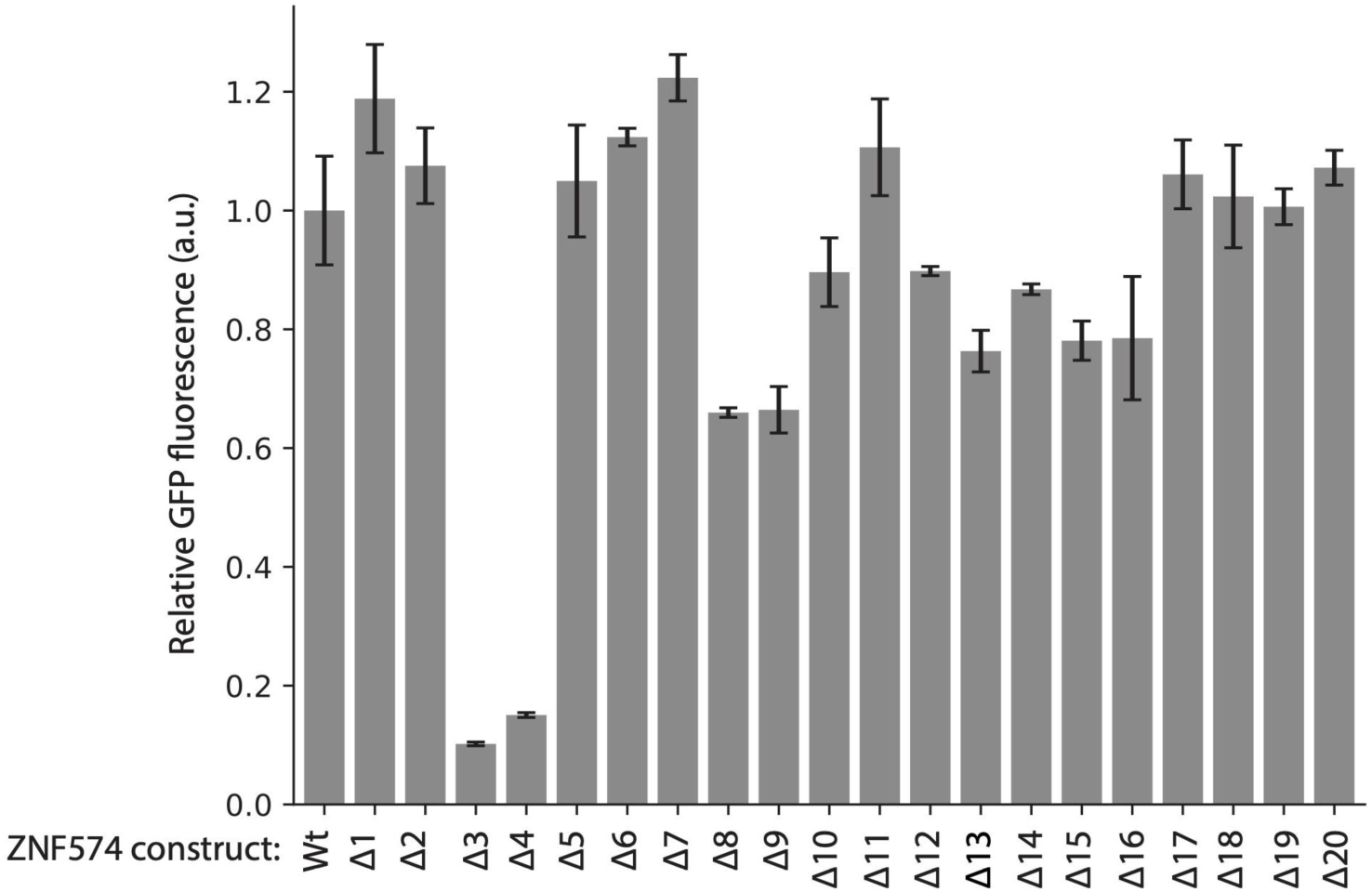
Expression levels of ZNF574 C2H2 deletes. GFP-tagged wild type ZNF574 or ZNF574 harboring a deletion of a single C2H2 domain were transduced in K562 cells and their relative GFP fluorescence was measured via flow cytometry (mean ± SD, *N=3*).

**SFig. 15.**
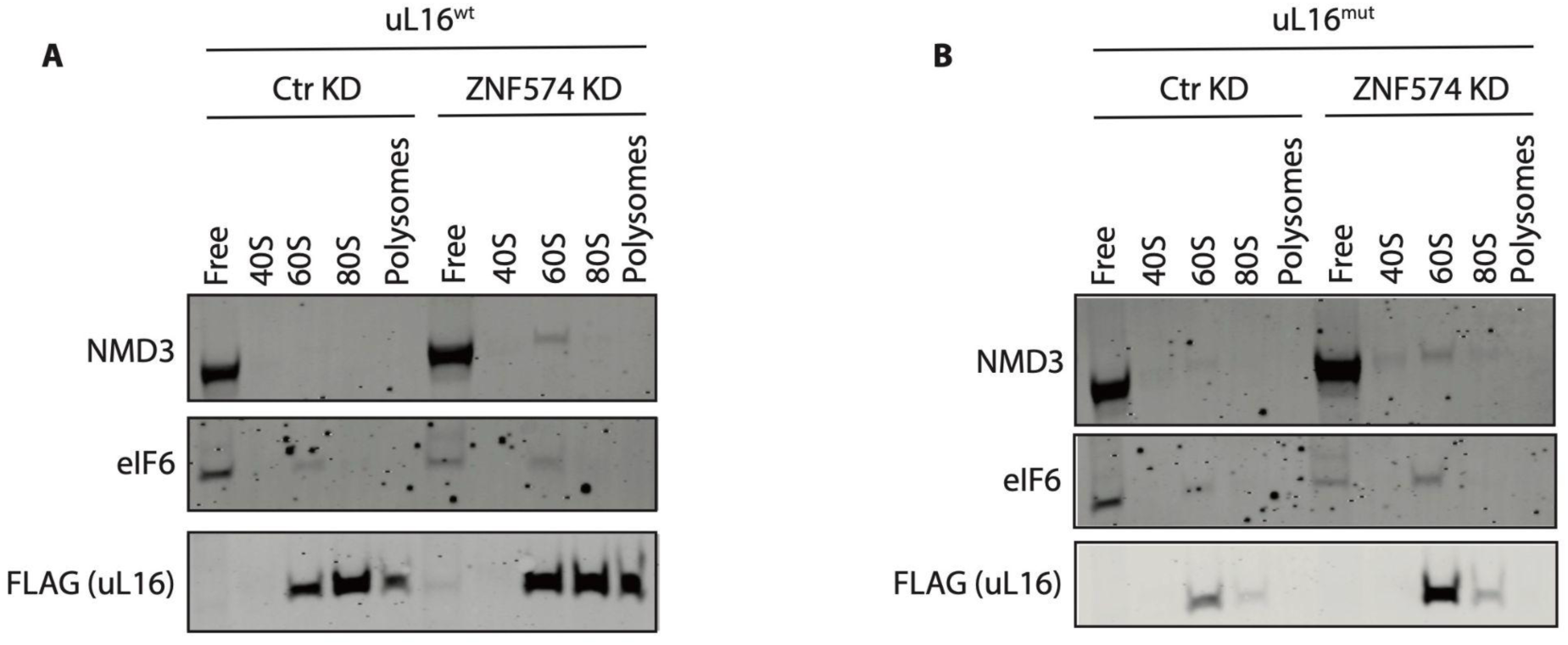
Knocking down ZNF574 traps ribosome biogenesis factors on defective large subunits. **A, B.** Cell lysates from K562 CRISPRi cells expressing wild type (A) or mutant (B) uL16-FLAG and control sgRNA or sgRNA targeting ZNF574 were analyzed on polysome gradients. Fractions corresponding to ribosome-free cytoplasm, 40S, 60S, 80S, and polysomes were collected and the presence of NMD3 and eIF6 in each fraction was detected via western blotting.

**SFig. 16.**
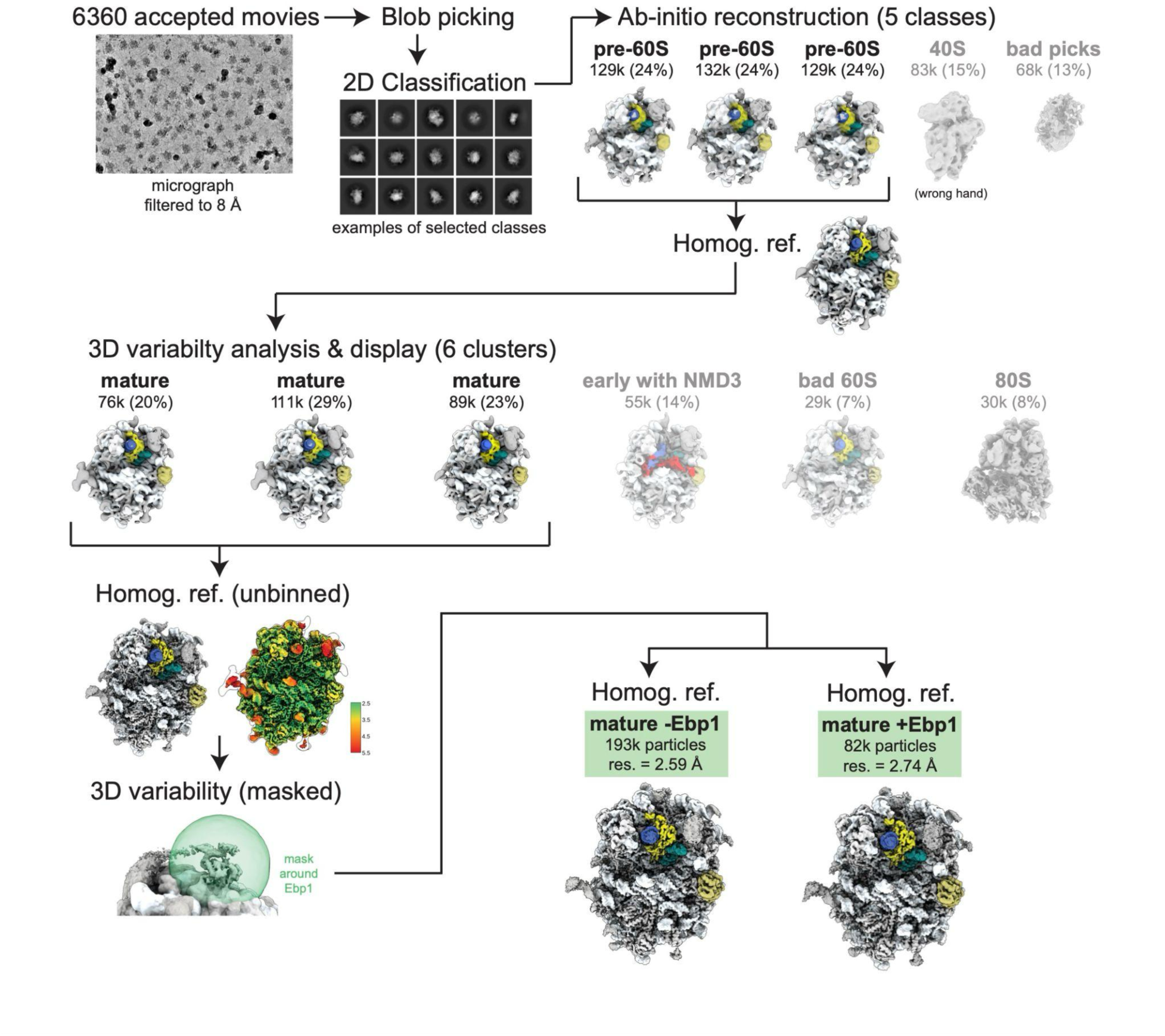
Processing scheme for cryo-EM data. Simplified diagram showing the steps of 3D classification and refinement resulting in the high-resolution reconstruction of human pre-60S incorporating uL16^mut^. Maps were colored according to the following scheme (CSS color names): alice blue (rRNA), light gray (protein), royal blue (helix 38), dark cyan (helix 89), yellow (uL16^mut^), dark khaki (eIF6), red (NMD3). Masks are represented as partially transparent green spheres.

