## Supplementary material for "ZNF574 is a Quality Control Factor For Defective Ribosome Biogenesis Intermediates": Suplemental_Materials: KEY RESOURCES TABLE_Akers.pdf

| REAGENT or RESOURCE | SOURCE | IDENTIFIER |
|---|---|---|
| **Antibodies** | | |
| Mouse anti-GFP | Invitrogen | RRID:AB_10979281 |
| Rabbit anti-NMD3 | Bethyl Laboratories, Inc. | RRID:AB_2782088 |
| Rabbit anti-EIF6 | Bethyl Laboratories, Inc. | RRID:AB_2780928 |
| Rat anti-HA | Roche | RRID:AB_390918 |
| Rabbit anti-RPL10 | Invitrogen | RRID:AB_2646721 |
| Rabbit anti-RPS24 | Abcam | RRID:AB_2714188 |
| Rabbit anti-RPL7 | Abcam | RRID:AB_1270391 |
| Mouse anti-FLAG | Sigma | RRID:AB_259529 |
| Mouse anti-tubulin | Sigma | RRID:AB_477579 |
| Rabbit anti-ZNF574 | Sigma | RRID:AB_2671105 |
| Mouse anti-Keima | MBL | M126-3M |
| **Chemicals, Peptides, and Recombinant Proteins** | | |
| Cycloheximide | Sigma-Aldrich | C4859-1ML |
| Bortezomib | Sigma | 5043140001 |
| Leupeptin Hemisulfate | Selleckchem | S7380 |
| Chloroquine diphosphate | Selleckchem | S4157 |
| Torin1 | Selleckchem | S2827 |
| Doxicyclyne monohydrate | Sigma | D1822 |
| **Deposited Data** | | |
| CRISPRi FACS screen gene-level table | This study | Table S1 |
| CRISPRi growth screen gene-level table | This study | Table S2 |
| uL16 IP-MS data | This study | Table S3, PXD051646 |
| ZNF598 IP-MS data | This study | Table S4, PXD051645 |
| **Experimental Models: Cell Lines** | | |
| K562 CRISPRi | (Gilbert et al., 2014) | N/A |
| K562 CRISPRi SFFV:uL16$^{wt}$-GFP | This study | N/A |
| K562 CRISPRi SFFV:uL16$^{mut}$-GFP | This study | N/A |
| K562 CRISPRi SFFV:uL16$^{wt}$-RFP | This study | N/A |
| K562 CRISPRi SFFV:uL16$^{mut}$-RFP | This study | N/A |
| K562 CRISPRi SFFV:uL16$^{wt}$-FLAG-IRES-mCherry | This study | N/A |
| K562 CRISPRi SFFV:uL16$^{mut}$-FLAG-IRES-mCherry | This study | N/A |
| K562 CRISPRi tet3G:uL16$^{wt}$-GFP | This study | N/A |
| K562 CRISPRi tet3G:uL16$^{wt}$-GFP sgGal4 | This study | |
| K562 CRISPRi tet3G:uL16$^{wt}$-GFP; sgGal4; SFFV:GFP-ZNF574(wt) | This study | N/A |

| | | |
|---|---|---|
| K562 CRISPRi tet3G:uL16$^{wt}$-GFP; sgGal4; SFFV:GFP-ZNF574(deltaC2H2#17-20) | This study | N/A |
| K562 CRISPRi tet3G:uL16$^{wt}$-GFP sgZNF574 | This study | N/A |
| K562 CRISPRi tet3G:uL16$^{wt}$-GFP; sgZNF574; SFFV:GFP-ZNF574(wt) | This study | N/A |
| K562 CRISPRi tet3G:uL16$^{wt}$-GFP; sgZNF574; SFFV:GFP-ZNF574(deltaC2H2#17-20) | This study | N/A |
| K562 CRISPRi tet3G:uL16$^{mut}$-GFP | This study | N/A |
| K562 CRISPRi tet3G:uL16$^{mut}$-GFP sgGal4 | This study | N/A |
| K562 CRISPRi tet3G:uL16$^{mut}$-GFP; sgGal4; SFFV:GFP-ZNF574(wt) | This study | N/A |
| K562 CRISPRi tet3G:uL16$^{mut}$-GFP; sgGal4; SFFV:GFP-ZNF574(deltaC2H2#17-20) | This study | N/A |
| K562 CRISPRi tet3G:uL16$^{mut}$-GFP sgZNF574 | This study | N/A |
| K562 CRISPRi tet3G:uL16$^{mut}$-GFP; sgZNF574; SFFV:GFP-ZNF574(wt) | This study | N/A |
| K562 CRISPRi tet3G:uL16$^{mut}$-GFP; sgZNF574; SFFV:GFP-ZNF574(deltaC2H2#17-20) | This study | N/A |
| K562 CRISPRi SFFV: GFP-ZNF574 | This study | N/A |
| K562 CRISPRi SFFV: GFP-ZNF574; SFFV:uL16$^{mut}$-RFP; sgZNF574 | This study | N/A |
| K562 CRISPRi SFFV: GFP-ZNF574deltaC2H2#1; SFFV:uL16$^{mut}$-RFP; sgZNF574 | This study | N/A |
| K562 CRISPRi SFFV: GFP-ZNF574deltaC2H2#2; SFFV:uL16$^{mut}$-RFP; sgZNF574 | This study | N/A |
| K562 CRISPRi SFFV: GFP-ZNF574deltaC2H2#3; SFFV:uL16$^{mut}$-RFP; sgZNF574 | This study | N/A |
| K562 CRISPRi SFFV: GFP-ZNF574deltaC2H2#4; SFFV:uL16$^{mut}$-RFP; sgZNF574 | This study | N/A |
| K562 CRISPRi SFFV: GFP-ZNF574deltaC2H2#5; SFFV:uL16$^{mut}$-RFP; sgZNF574 | This study | N/A |
| K562 CRISPRi SFFV: GFP-ZNF574deltaC2H2#6; SFFV:uL16$^{mut}$-RFP; sgZNF574 | This study | N/A |
| K562 CRISPRi SFFV: GFP-ZNF574deltaC2H2#7; SFFV:uL16$^{mut}$-RFP; sgZNF574 | This study | N/A |
| K562 CRISPRi SFFV: GFP-ZNF574deltaC2H2#8; SFFV:uL16$^{mut}$-RFP; sgZNF574 | This study | N/A |
| K562 CRISPRi SFFV: GFP-ZNF574deltaC2H2#9; SFFV:uL16$^{mut}$-RFP; sgZNF574 | This study | N/A |
| K562 CRISPRi SFFV: GFP-ZNF574deltaC2H2#10; SFFV:uL16$^{mut}$-RFP; sgZNF574 | This study | N/A |

| | | |
|---|---|---|
| K562 CRISPRi SFFV: GFP-ZNF574deltaC2H2#11; SFFV:uL16$^{mut}$-RFP; sgZNF574 | This study | N/A |
| K562 CRISPRi SFFV: GFP-ZNF574deltaC2H2#12; SFFV:uL16$^{mut}$-RFP; sgZNF574 | This study | N/A |
| K562 CRISPRi SFFV: GFP-ZNF574deltaC2H2#13; SFFV:uL16$^{mut}$-RFP; sgZNF574 | This study | N/A |
| K562 CRISPRi SFFV: GFP-ZNF574deltaC2H2#14; SFFV:uL16$^{mut}$-RFP; sgZNF574 | This study | N/A |
| K562 CRISPRi SFFV: GFP-ZNF574deltaC2H2#15; SFFV:uL16$^{mut}$-RFP; sgZNF574 | This study | N/A |
| K562 CRISPRi SFFV: GFP-ZNF574deltaC2H2#16; SFFV:uL16$^{mut}$-RFP; sgZNF574 | This study | N/A |
| K562 CRISPRi SFFV: GFP-ZNF574deltaC2H2#17; SFFV:uL16$^{mut}$-RFP; sgZNF574 | This study | N/A |
| K562 CRISPRi SFFV: GFP-ZNF574deltaC2H2#18; SFFV:uL16$^{mut}$-RFP; sgZNF574 | This study | N/A |
| K562 CRISPRi SFFV: GFP-ZNF574deltaC2H2#19; SFFV:uL16$^{mut}$-RFP; sgZNF574 | This study | N/A |
| K562 CRISPRi SFFV: GFP-ZNF574deltaC2H2#20; SFFV:uL16$^{mut}$-RFP; sgZNF574 | This study | N/A |
| HEK293T SFFV: GFP-ZNF574 | This study | N/A |
| U2OS CRISPRi SFFV:uL16$^{wt}$-FLAG-IRES-mCherry; sgGal4 | This study | N/A |
| U2OS CRISPRi SFFV:uL16$^{wt}$-FLAG-IRES-mCherry; sgZNF574 | This study | N/A |
| U2OS CRISPRi SFFV:uL16$^{mut}$-FLAG-IRES-mCherry; sgGal4 | This study | N/A |
| U2OS CRISPRi SFFV:uL16$^{mut}$-FLAG-IRES-mCherry; sgZNF574 | This study | N/A |
| **Oligonucleotides** | | |
| See Table S5 for qPCR primers | This study | N/A |
| See Table S6 for protospacer sequences | This study | N/A |
| **Recombinant DNA** | | |
| hCRISPRi- v2 library | Horlbeck et al., 2016 | Addgene, Cat#83969 |
| hCRISPRi_dual_1_2 pooled library | Replogle et al., 2022 | Addgene, Cat#187246 |
| See Table S7 for plasmids | This study | N/A |
| pU6-sgRNA EF1alpha-puro-T2A-BFP | Gilbert et al., 2014 | Addgene Cat#60955 |
| **Software and Algorithms** | | |
| Screen Processing Pipeline | Horlbeck et al., 2016 | https://github.com/mhorlbeck/ScreenProcessing |

| | | |
|---|---|---|
| FlowJo 8.8.6 | FlowJo | https://www.flowjo.com |
| MAGeCK | Li et al., 2014 | https://hpc.nih.gov/apps/MAGeCK.html |
| Fiji | Schindelin et al., 2012 | https://fiji.sc/ |

**Cell culture, viral production, and construction of reporter cell lines**

K562 cells were grown in RPMI-1640 with 25 mM HEPES, 2.0 g/L NaHCO3, 0.3 g/L L-Glutamine supplemented with 10% FBS, 2 mM glutamine, 100 units/mL penicillin and 100 µg/mL streptomycin. HEK293T and U2OS cells were grown in Dulbecco's modified eagle medium (DMEM) in 10% FBS, 2 mM glutamine, 100 units/mL penicillin and 100 µg/mL streptomycin. All cell lines were obtained from ATCC and were tested for Mycoplasma quarterly.

**Generation of cell lines**

To generate the K562 cell line stably expressing dCas9-KRAB, cells were stably transduced with a lentiviral vector expressing dCas9-BFP-KRAB from an EF1-α promoter with an upstream ubiquitous chromatin opening element (SFFV-dCas9-BFP-KRAB, Addgene #85969) and selected for BFP-positive cells using two rounds of fluorescence-activated cell sorting (FACS) on a BD FACSAria2.

Reporter cell lines (uL16 reporters, ZNF574 reporters) were stably transduced with the corresponding lentiviral vectors (Table S5), and positive cells were isolated by FACS on a BD FACSAria3.

Individual gene knockdowns were carried out by selecting sgRNA protospacers from the compact hCRISPRi-v2 library and cloning into lentiviral plasmid pU6-sgRNA EF1α-puro-t2a-BFP (Addgene 60955) as previously described[1]. Protospacer sequences used for individual knockdowns are listed in Table S6. The resulting sgRNA expression vectors were packaged into lentivirus by transfecting HEK293T with standard packaging vectors using PEI transfection. The viral supernatant was harvested 2–3 days after transfection and filtered through 0.45 µm PVDF filter and/or frozen prior to transduction into CRISPRi knockdown cell lines described above.

**PEI transfection**

Linear PEI at 25 kDa (Polysciences cat #23966-2) was dissolved in PBS pH 4.5 to 1 mg/ml. DNA and PEI were mixed in serum-free media at 1:3 ratio and incubated at room temperature for 15 minutes. Transfection mix was added to HEK293T cells and media was changed after 24 hours.

**Genome-scale CRISPRi screening**

Genome-scale screens were conducted similarly to previously described screens[1–3].

*Growth based screen*

The CRISPRi compact library (5 sgRNA/TSS) CRISPRi-v2 (Addgene, Cat#83969) was transduced in duplicate into K562 CRISPRi cells stably expressing $uL16^{wt}$ or $uL16^{mut}$ at MOI < 1 (percentage of transduced cells 2 days after transduction: 20%–40%). Replicates were maintained separately in 1 L of RPMI-1640 in spinner flasks for the course of the screen. 2 days after transduction, the cells were selected with 1 mg/mL puromycin for 3 days, at which point transduced cells accounted for 80%–95% of the population. Cells were allowed to recover to > 80% cell viability, as measured on a Cell Countess (Invitrogen). Cells were maintained in spinner flasks by daily dilution to $0.5x10^6$ cells /mL at an average coverage of greater than 1000 cells per sgRNA for the duration of the screen. Cells were harvested after 10 cell doublings. Genomic DNA was isolated from frozen cells using Macherey-Nagel NucleoSpin L kit (cat: 740954.20), and the sgRNA-encoded regions were enriched, amplified, and prepared for sequencing as described previously[2]. Sequencing reads were aligned to the CRISPRi v2 library sequences, counted, and quantified using the Python-based Screen Processing pipeline[1] (https://github.com/mhorlbeck/ScreenProcessing). Calculation of phenotypes and Mann-Whitney p-values was performed as described previously[1,2]. Phenotypes from sgRNAs targeting the same gene were collapsed into a single stabilization phenotype using the average of the top three scoring sgRNAs (by absolute value) and assigned a p-value using the Mann-Whitney test of all sgRNAs targeting the same gene compared to the non-targeting controls. All additional CRISPR screen data analyses were performed in Python 2.7 and Python 3 using a combination of Numpy (v1.12.1), Pandas (v0.17.1), and Scipy (v0.17.0). Gene-level phenotypes are available in Table S1.

*FACS based screen*

The CRISPRi compact library (hCRISPRi_dual_1_2 pooled library, Addgene, Cat#187246) was transduced in duplicate into K562 CRISPRi cells stably expressing $uL16^{mut}$-RFP at MOI < 1 (percentage of transduced cells 2 days after transduction: 20%–40%). Replicates were maintained separately in four T125 flasks per replica for the course of the screen. 2 days after transduction, the cells were selected with 1 mg/mL puromycin for 3 days, at which point transduced cells accounted for 80%–95% of the population. Cells were allowed to recover to > 80% cell viability, as measured on a Cell Countess (Invitrogen). The cells were maintained in T125 flasks by daily dilution to $0.5x10^6$ cells /mL at an average coverage of greater than 1000 cells per sgRNA for the duration of the screen. Cells were sorted using BD FACS Aria3 two days after recovery based on RFP fluorescence of $uL16^{mut}$-RFP reporter. Cells with the highest (~30%) and lowest (~30%) RFP expression were collected and snap-frozen. Approximately 10 million cells were collected per bin. Genomic DNA was isolated from frozen cells using Macherey-Nagel NucleoSpin L kit (cat: 740954.20). The sgRNA-encoded regions were enriched, amplified, and prepared for sequencing as described previously[3]. Sequencing reads were aligned to the CRISPRi compact library sequences, counted, and quantified using the MAGeCK pipeline[4]. RFP stability phenotypes were calculated by $\log_2$ transforming the ratio of the sgRNA counts in the top 30% sorted samples and the sgRNA counts in the bottom 30%. All additional CRISPR screen data analyses were performed in Python 3 using a combination of Numpy (v1.12.1), Pandas (v0.17.1), and Scipy (v0.17.0). Gene-level phenotypes are available in Table S2.

**Individual evaluation of sgRNA phenotypes**

For individual evaluation and re-testing of sgRNA phenotypes, sgRNA protospacers were individually cloned by annealing complementary synthetic oligonucleotide pairs (Integrated DNA Technologies) with flanking BstXI and

BlpI restriction sites and ligating the resulting double-stranded segment into BstXI/BlpI-digested pCRISPRi-v2. Protospacer sequences used for individual evaluation are listed in Table S6. The resulting sgRNA expression vectors were individually packaged into lentivirus. These sgRNA vectors were transduced into K562-dCas9 cells expressing uL16 reporters tagged with RFP or GFP at MOI < 1 (20 – 40% infected cells). RFP and GFP protein levels were measured by flow cytometry at day 5 post infection. Mean fluorescent values were calculated for each cell line using FlowJo software and compared to uninfected levels.

**Internally controlled growth assays**

For uL16 growth assay (Fig 2B), K562 cells were transduced with uL16$^{wt}$-RFP or uL16$^{mut}$-RFP expression constructs resulting in 30 – 50% infected cells. 48 hours post infection was considered time point zero and the ratio of RFP positive to negative cells was normalized to this time point. Cells were maintained at $0.5 \times 10^6$ cells/ml for the duration of the experiment. Cells were grown for 10 days and the ratio of RFP positive to negative cells was measured daily on an Attune NxT flow cytometer (Thermo Fisher Scientific).

For sgRNA knockdown experiments (Fig. 2D), K562 cells expressing uL16$^{wt}$-RFP or uL16$^{mut}$-RFP were transduced with sgRNA expression constructs (control or targeting ZNF574) resulting in 30 – 50% infected cells. 48 hours post infection is considered time point zero and the ratio of BFP positive to negative cells is normalized to this time point. Cells were maintained at $0.5 \times 10^6$ cells/ml for the duration of the experiment. Cells were grown for 10 days and the ratio of BFP positive to negative cells was measured daily on Attune NxT flow cytometer (Thermo Fisher Scientific).

**RT-qPCR:**

Total RNA was isolated using Zymo mini RNA prep (R2053). Reverse-transcription was carried out using M-MLV (Thermo Fisher Scientific) with random hexamer primers (Thermo Fisher Scientific, SO124) in the presence of RNaseIN Recombinant Ribonuclease Inhibitor (Thermo Fisher Scientific). Quantitative PCR (qPCR) was performed with SsoAdvanced Universal SYBR Green Supermix (Bio-Rad), according to the manufacturer's instructions on a CFX384 Touch Real-Time PCR Detection System (Bio-Rad). Experiments were performed in technical triplicates. RT-qPCR primers used are listed in Table S7. *GAPDH* was used as a housekeeping gene. Ectopic *uL16* was detected via primers targeting the GFP or RFP tag, and the endogenous *uL16* was detected via primers targeting the endogenous *uL16* 3'UTR, which was not part of the ectopically expressed *uL16* constructs. The relative fold gene expression was calculated using the delta-delta Ct method[5].

**Polysome gradients:**

Sucrose density gradients (10%–50%) were prepared and measured in Seaton Open Top Polyclear centrifuge tubes (Thermo Fisher Scientific, NC9863486) using a BioComp Gradient Station (BioComp Instruments) according to the manufacturer's instructions. Sucrose solutions were prepared in 20 mM Tris pH 7.5, 150 mM NaCl, 5 mM $MgCl_2$, 1 mM DTT buffer. Samples were lysed in 20 mM Tris pH 7.5, 150 mM NaCl, 5 mM $MgCl_2$, 1% Triton x-100, 1 mM DTT, 24 U/ml Turbo DNase (Ambion), 20 U/mL SUPERaseIn (Ambion) and loaded onto gradients, which were spun for 1 hour 45 minutes at 40,000 rpm, 4°C in a SW41 rotor (Beckman Coulter). Samples were finally loaded onto Biocomp Gradient Station and the 260 nm or GFP absorbance was read using a Triax™ flow cell (BioComp Instruments).

**Proteasome inhibition**

Bortezomib (Sigma 5043140001) was dissolved in DMSO and added to cells at 50 nM final concentration. Cells were harvested after 5 hours and analyzed for GFP or RFP fluorescence via Attune NxT flow cytometer (Thermo Fisher Scientific) or lysed for Western blot.

**HA-Ubiquitin assay**

293T cells were transfected with HA-tagged ubiquitin plasmid using TransIT-LTI Transfection Reagent (Mirus, MIR 2306). The proteasomal inhibitor bortezomib was added at 50 nM final concentration 24 hours after transfection, and cells were harvested for experiments 5 hours post proteasomal inhibition.

**Immunoprecipitation (IP) and western blot:**

Cells were lysed in buffer containing 50 mM HEPES, 100 mM KCl, 15 mM $Mg(OAc)_2$, 5% glycerol, 0.25% NP40, cOmplete™ Mini protease inhibitor cocktail, EDTA-free (Roche), 20U/ml SUPERaseIn™ RNase Inhibitor (Thermo), 24 U/ml TURBO™ DNase (Thermo). If the lysate was used for ubiquitin detection, the lysis buffer and all wash buffers were also supplemented with 5 mM N-ethylmaleimide (NEM) (Sigma, E3876-5G). The lysates were cleared by centrifugation at 8000 x g for 10 min and bound to FLAG magnetic beads (Millipore M8823-1ML) for 1h or overnight at 4°C. Beads were washed 3 times with Wash Buffer 1 (50 mM HEPES, 100 mM KCl, 15 mM $Mg(OAc)_2$, 5% glycerol, 0.1% NP40), and 3 times with Wash Buffer 2 (50 mM HEPES, 100 mM KCl, 15 mM $Mg(OAc)_2$, 0.05% NP40). Bound material was competitively eluted with FLAG peptide (0.5 mg/ml FLAG peptide (Sigma F4799) in 25 mM Tris 7.5, 150 mM NaCl). Relevant IP fractions were boiled in Laemmli Buffer for 10 min at 90°C.

Proteins were separated on Bolt® 4-12% Bis-tris gels (Thermo Fisher Scientific), transferred to PVDF membrane using the Mini Trans-Blot Cell (Bio-Rad) according to the manufacturer's instructions, blocked with 5% milk in TBS, and subsequently probed. LI-COR IRDye700 anti-mouse (LI-COR, 926-68070), LI-COR IRDye800 anti-rabbit (LI-COR, 926-32211), and LI-COR IRDye800 anti-rat (LI-COR, 926-32219) secondary antibodies were then used at 1:10,000 dilution. All blots were visualized using the LI-COR Odyssey system.

**Microscopy**

U2OS cells expressing uL16 reporters and sgRNAs were plated in 8-well ibiTreat µSlide (ibidi 80826) at 20–25,000 cells/well. On the following day, cells were fixed with 4% PFA for 10 min at room temperature, washed with PBS (137 mM NaCl, 2.7 mM KCl, 10 mM Na2HPO4, 1.8 mM K2HPO4), and permeabilized with 0.5% Triton in PBS for 10 min at room temperature. Slides were then treated with blocking buffer (5% goat serum (Jackson Immuno Research) in PBS) for 1 hour at room temperature. Antibodies were diluted in a blocking buffer and incubated with cells at 4°C overnight. After three washes with PBST (1x PBS, 0.1% Tween), cells were incubated with secondary antibodies conjugated to Alexa 488 or Alexa 568, and DAPI (1 ug/ml, Sigma D9542) for 1 hour at room temperature. Slides were imaged on a Nikon Eclipse E800 microscope. Images were analyzed using Fiji[6].

**Mass Spectrometry**

*LC-MS/MS Analysis*

Sliced gels were subjected to in-gel digestion and desalted using ziptips. Peptides were analyzed by liquid chromatography–tandem mass spectrometry (LC-MS) on an EasyLC1200 system (Thermo) connected to a high-performance quadrupole Orbitrap Q Exactive (Thermo). Peptides were separated using ES803 (50 cm) column. Peptides were eluted from the analytical column by a gradient from 3 to 5% solvent B (80% (v/v) acetonitrile 0.1% (v/v)) over 1 minute, 5 to 28% solvent B over 105 minutes, and from 28 to 44% solvent B over 15 minutes, followed by a short wash (15 min) at 90% solvent B. Precursor scan was from mass-to-charge ratio (m/z) 375 to 1600 (resolution 120,000; AGC 3.0 $e^6$, maximum injection time 100 ms) and top 20 most intense multiply charged precursors were selected for fragmentation (resolution 15,000, AGC 5E4, maximum injection time 60ms, isolation window 1.0 m/z, minimum AGC target 1.2 $e^3$, intensity threshold 2.0 $e^4$, include charge state = 2 – 8). Peptides were fragmented with higher-energy collision dissociation (HCD) with normalized collision energy (NCE) 27. Dynamic exclusion was enabled for 24 seconds.

*Processing of Mass Spectrometry Data*

Mass spectrometry data was searched using FragPipe/MSFragger for peptide identification and Label-Free Quantitation (LFQ). Raw MS/MS data was loaded into the FragPipe interface and searched and quantified with the MSFragger search engine (v20.0)[7], using default settings and allowing normalization and match between runs. Data was searched against the SwissProt *Homo Sapiens* protein database, concatenated with decoy protein sequences (84188 total entries). A 10 ppm precursor mass tolerance and 20 ppm MS/MS2 tolerance were allowed. Carbamidomethyl (cysteine (C)) was searched as a constant modification. Variable modifications allowed include protein N-terminal acetylation, methionine (M) oxidation, N-terminal acetylation with M oxidation, N-terminal M-loss, N-terminal M-loss with acetylation, and peptide N-terminal Gln conversion to pyroglutamate, with two maximum variable modifications per peptide. Cleavage specificity was set to trypsin, allowing one missed cleavage. For all samples, a 5% FDR was permitted for protein, and a 1% FDR was permitted for peptide identifications. LFQ was run on default settings, using match between runs.

*Statistics processing using Perseus*

The search results were analyzed separately in Perseus (version 2.0.10.0). MSFragger's 'combined_proteins' were loaded in Perseus, with LFQ intensities specified as 'Main' columns. Decoys and/or contaminants were removed, then the MaxLFQ intensities were log2 transformed and quality checked with histograms. Non-reproducible identifications were removed (keep only rows with at least 3 valid values in either ZNF574-GFP replicates or GFP replicates), then missing values were imputed from a normal distribution downshifted by 1.8 and a width of 0.3

column-wise. A two-sided T-test with S0=0.5 and FDR=0.05 was used to determine the significance and enrichment of each identified peptide or protein.

### Cryo-EM

*Sample preparation*

K562 cells expressing $uL16^{mut}$ and sgRNA targeting ZNF574 were lysed in Lysis Buffer (20 mM Tris pH 7.5, 150 mM NaCl, 5 mM $MgCl_2$, 1% Triton x-100, 1 mM DTT, 24 U/ml Turbo DNase (Ambion), 20 U/mL SUPERaseIn (Ambion), 1x Halt$^{TM}$ Protease Inhibitor (Thermo, PI78429), 1 U/ml Apyrase (NEB, M0398S)). Lysate was clarified by two centrifugations at 20, 000 rpm in Ti70 rotor (Beckman Coulter) for 20 min. A discontinuous sucrose gradient was prepared by layering 6 ml 25% sucrose above 6 ml 48% sucrose dissolved in 20 mM Tris pH 7.5, 100 mM NaCl, 6 mM $Mg(OAc)_2$, 2 mM DTT. Ribosomes were pelleted through the discontinuous gradient for 20 h at 26, 000 rpm. The ribosome pellet was resuspended in 50 mM HEPES pH 7.6, 6 mM $Mg(OAc)_2$, 150 mM KCl, 6.8% sucrose (w/v), 2 mM DTT. Sucrose density gradients (10%–50%) were prepared and measured in Seaton Open Top Polyclear centrifuge tubes (Thermo Fisher Scientific, NC9863486) using a BioComp Gradient Station (BioComp Instruments) according to the manufacturer's instructions. Sucrose solutions were prepared in 20 mM Tris pH 7.5, 150 mM NaCl, 5 mM $MgCl_2$, and 1 mM DTT buffer. Resuspended ribosomes were loaded onto gradients and spun for 1 hour 45 minutes at 40,000 rpm, 4°C in a SW41 rotor (Beckman Coulter). Samples were finally loaded onto Biocomp Gradient Station and the 260 nm absorbance was read using a Triax$^{TM}$ flow cell (BioComp Instruments). 60S ribosome fractions were manually collected and buffer exchanged into 20 mM HEPES pH7.6, 5 mM MgCl2, 100 mM KCl, 2 mM DTT using Zeba$^{TM}$ Spin Desalting Columns 7K MWCO (Thermo). Finally, samples were concentrated using Amicon$^{TM}$ Ultra-15 Centrifugal Filter Unit 100K MWCO (Millipore).

*Cryo-electron microscopy*

Quantifoil R2/2 holey carbon copper grids (Quantifoil Micro Tools) were covered with a homemade sheet of carbon with a thickness of 1 nm. Grids were glow-discharged using an easiGlow Discharge cleaning system (Pelco) for 15 s with a current of 15 mA. Using a Vitrobot (Thermo Fisher Scientific) with chamber temperature set to 4 °C and chamber humidity set to 100%, a 5 µl sample of 80 nM purified 60S ribosomal subunits was applied. After 30 s of incubation, the excess sample was blotted for 1–6 s using a blot force of 15, and the grid was plunge-frozen in a mixture of ethane and propane (1:2 ratio).

For electron microscopy, grids were loaded into a Titan Krios (ThermoFisher Scientific) cryo-transmission electron microscope with an operating voltage of 300 kV. Data was collected in counting and super-resolution mode using a K3 direct electron camera (Gatan) mounted to a GIF Quantum LS energy filter (Gatan). The magnification of the microscope was set to 81,000× which resulted in a pixel size of 1.06 Å/pixel. The data collection was controlled with the EPU program (ThermoFisher Scientific). 40 frames were recorded per movie, with a defocus range between -1 and -2.5 µm (0.3 µm increments), and a total dose of 60 $e^-/Å^2$. The energy filter slit width was set to 20 eV.

*Cryo-EM data processing*

A total of 7761 movies were imported into CryoSPARC Live[8] to perform motion correction, CTF estimation and particle picking. Exposures were rejected based on ice thickness and decofus range, resulting in 6360 accepted movies. Particles were picked with a circular blob using a particle diameter between 250–350 Å and a minimal separation distance of 0.5 diameters. Particles were extracted using a box size of 560 pixels and fourier-cropped to 186 pixels for initial processing.

2D classification was performed in CryoSPARC Live using 100 classes. 38 classes were showing ribosome-like 2D averages and were selected for ab-initio reconstruction in CryoSPARC using 5 classes. The three 60S-like classes were used for homogeneous refinement and subsequently subjected to global 3D variability analysis, which was displayed with 6 clusters, three of which corresponded to mature pre-60S and one of which corresponded to an NMD3-bound assembly intermediate.

Particles from clusters representing mature pre-60S were extracted using a box size of 560 pixels and refined to a resolution of 2.5 Å using homogeneous refinement and further classified into particles with and without Ebp1 by a masked 3D variability analysis. The class without Ebp1 was homogeneously refined to 2.59 Å.

A simplified scheme of the processing workflow is shown in Fig. S16.

*Model building and visualization*

A three-dimensional structure model was built using a deposited PDB structure as a starting model. We used Coot 0.9.8.5[9] for structure building, editing and real-space refinement. As a starting model for building of the 60S subunit we used a recent high-resolution structure (PDB accession number 8A3D)[10], and re-built the P-site loop of uL16 to account for the deletion of residues 102–111. We added the missing modifications of the ribosomal RNA nucleotides according to published results of quantitative mass spectrometry[11], using eLBOW[12] to generate the restraint file for 2-O-methyl-pseudouridine based on the .cif file downloaded from the protein database (https://www.rcsb.org/ligand/UY1). Additionally, we modeled additional well-resolved parts of the structure by hand-building. We docked the helix of ZNF622 that is resolved within the peptide exit tunnel, from a published model (PDB: 6LSR)[13]. The structure was refined using Phenix[14].

CryoSPARC was used to generate low-pass filtered maps. Figures were generated with UCSF ChimeraX. Labels and schemes were added in Adobe Illustrator.

**Table S1: Gene level growth CRISPRi**

**Table S2: Gene level FACS CRISPRi**

**Table S3: MS for uL16**

**Table S4: MS for ZNF574**

**Table S5: Plasmids used in this study**

**Table S6: Protospacers used in this study**

**Table S7: RT-qPCR primers used in this study**

**Table S8: Statistics of cryo-EM data collection and refinement.**

**Supplemental Material Bibliography:**


1. Horlbeck, M.A., Gilbert, L.A., Villalta, J.E., Adamson, B., Pak, R.A., Chen, Y., Fields, A.P., Park, C.Y., Corn, J.E., Kampmann, M., et al. (2016). Compact and highly active next-generation libraries for CRISPR-mediated gene repression and activation. Elife *5*, e19760. 10.7554/eLife.19760.
2. Gilbert, L.A., Horlbeck, M.A., Adamson, B., Villalta, J.E., Chen, Y., Whitehead, E.H., Guimaraes, C., Panning, B., Ploegh, H.L., Bassik, M.C., et al. (2014). Genome-Scale CRISPR-Mediated Control of Gene Repression and Activation. Cell *159*, 647–661. 10.1016/J.CELL.2014.09.029.
3. Replogle, J.M., Bonnar, J.L., Pogson, A.N., Liem, C.R., Maier, N.K., Ding, Y., Russell, B.J., Wang, X., Leng, K., Guna, A., et al. (2022). Maximizing CRISPRi efficacy and accessibility with dual-sgRNA libraries and optimal effectors. Elife *11*. 10.7554/ELIFE.81856.
4. Li, W., Xu, H., Xiao, T., Cong, L., Love, M.I., Zhang, F., Irizarry, R.A., Liu, J.S., Brown, M., and Liu, X.S. (2014). MAGeCK enables robust identification of essential genes from genome-scale CRISPR/Cas9 knockout screens. Genome Biol *15*, 554. 10.1186/S13059-014-0554-4.
5. Livak, K.J., and Schmittgen, T.D. (2001). Analysis of relative gene expression data using real-time quantitative PCR and the 2(-Delta Delta C(T)) Method. Methods *25*, 402–408. 10.1006/METH.2001.1262.
6. Schindelin, J., Arganda-Carreras, I., Frise, E., Kaynig, V., Longair, M., Pietzsch, T., Preibisch, S., Rueden, C., Saalfeld, S., Schmid, B., et al. (2012). Fiji: an open-source platform for biological-image analysis. Nature Methods 2012 9:7 *9*, 676–682. 10.1038/nmeth.2019.
7. Kong, A.T., Leprevost, F. V., Avtonomov, D.M., Mellacheruvu, D., and Nesvizhskii, A.I. (2017). MSFragger: ultrafast and comprehensive peptide identification in mass spectrometry–based proteomics. Nature Methods 2017 14:5 *14*, 513–520. 10.1038/nmeth.4256.
8. Punjani, A., Rubinstein, J.L., Fleet, D.J., and Brubaker, M.A. (2017). cryoSPARC: algorithms for rapid unsupervised cryo-EM structure determination. Nat Methods *14*, 290–296. 10.1038/NMETH.4169.
9. Emsley, P., Lohkamp, B., Scott, W.G., and Cowtan, K. (2010). Features and development of Coot. Acta Crystallogr D Biol Crystallogr *66*, 486. 10.1107/S0907444910007493.
10. Faille, A., Dent, K.C., Pellegrino, S., Jaako, P., and Warren, A.J. (2023). The chemical landscape of the human ribosome at 1.67 Å resolution. bioRxiv, 2023.02.28.530191. 10.1101/2023.02.28.530191.
11. Taoka, M., Nobe, Y., Yamaki, Y., Sato, K., Ishikawa, H., Izumikawa, K., Yamauchi, Y., Hirota, K., Nakayama, H., Takahashi, N., et al. (2018). Landscape of the complete RNA chemical modifications in the human 80S ribosome. Nucleic Acids Res *46*, 9289–9298. 10.1093/NAR/GKY811.
12. Moriarty, N.W., Grosse-Kunstleve, R.W., and Adams, P.D. (2009). electronic Ligand Builder and Optimization Workbench (eLBOW): a tool for ligand coordinate and restraint generation. Acta Crystallogr D Biol Crystallogr *65*, 1074–1080. 10.1107/S0907444909029436.
13. Liang, X., Zuo, M.Q., Zhang, Y., Li, N., Ma, C., Dong, M.Q., and Gao, N. (2020). Structural snapshots of human pre-60S ribosomal particles before and after nuclear export. Nat Commun *11*. 10.1038/S41467-020-17237-X.
14. Liebschner, D., Afonine, P. V., Baker, M.L., Bunkoczi, G., Chen, V.B., Croll, T.I., Hintze, B., Hung, L.W., Jain, S., McCoy, A.J., et al. (2019). Macromolecular structure determination using X-rays, neutrons and electrons: recent developments in Phenix. Acta Crystallogr D Struct Biol *75*, 861–877. 10.1107/S2059798319011471.
